# An open field phenomics resource for multimodal maize yield prediction across divergent environments

**DOI:** 10.64898/2026.09.17.752474

**Authors:** Aaron J. DeSalvio, Peiman Mohseni, Alper Adak, Seth C. Murray, Mustafa A. Arik, Raymond K. W. Wong, Jinha Jung, Dayane C. Lima, Alejandro C. Aviles, Edward Buckler, Nick Duffield, Jode Edwards, David Ertl, Sherry Flint-Garcia, Michael A. Gore, Candice N. Hirsch, James Holland, Shawn M. Kaeppler, Jarrod Miller, Maria Cinta Romay, James C. Schnable, Maninder P. Singh, Erin E. Sparks, Addie Thompson, Jacob D. Washburn, Teclemariam Weldekidan, Noah D. Winans, Natalia de Leon

**Author notes:** Contributing authors.

## Abstract

Temporal drone phenotyping captures crop development, but irregular flight schedules complicate comparisons across environments. We release curated imagery from 356 flights across 19 Genomes to Fields environments containing 1,180 maize (*Zea mays* L.) hybrids. To evaluate its utility, we integrated functional principal components of vegetation index and weather trajectories with genomic information. Combinined genomic and phenomic kernels improved yield prediction, reaching correlations up to *r* = 0.501 for held-out hybrids in environments represented in training and 0.408 when environments were also withheld. Accumulated growing degree days offered no consistent predictive advantage over days after planting, and weather contributed modest, task-dependent gains. A transformer neural process learned directly from irregular observations, serving as a novel application of neural process models in agriculture. Mapping vegetation index functional principal components identified recurrent quantitative trait loci on chromosomes 3 and 7. This resource and its reproducible analyses guide the use of temporal spectral data for crop prediction and genetic discovery.

## 1 Introduction

Understanding variability of quantitative terminal traits such as grain yield across production environments involves predicting within-environment genetic effects and genotype-by-environment (G×E) interactions. Maize (*Zea mays* L.) is cultivated on six continents and subject to intense artificial selection. While G×E interactions have diminished in modern temperate cultivars, they still comprise a substantial amount of variation that is resource intensive to measure [1].

Decades of genotyping advancements have allowed genomic selection, genomic prediction, and genome-wide association studies to link genotypes with phenotypes [2]. However, genomic markers are time-invariant, while crop development reflects dynamic environmental and G×E interactions. Consequently, genomics alone cannot fully capture the temporal processes in each environment underlying crop performance.

Weather measurements and data from remote sensing platforms, such as drones, provide complementary temporal information. Weather data characterize the environmental drivers of plant responses [3–5]. High-throughput phenotyping (HTP) and phenomics approaches use repeated spectral measurements to capture data related to plant responses to environmental conditions at whole plot or single plant resolution [6, 7]. Multimodal predictive approaches integrate dynamic enviromic and phenomic data alongside genomic marker information [8]. Statistical kernel and reaction norm models have employed genomic relationships, environmental factors, and crop growth model outputs to quantify environmental similarity and G×E [9–12]. Weather-based relationship matrices model environmental similarity, quantifying relatedness between environments as opposed to assuming they are unrelated [4, 13, 14].

Phenomic measurements, such as spectral imaging and near-infrared spectroscopy, have improved prediction of yield and other complex traits [15–19]. Machine and deep learning approaches serve as alternatives to kernel-based methods by learning latent representations across data modalities. Previous studies synthesizing enviromic and genomic predictors have used gradient boosting, employed transfer learning across traits and species, and integrated high-dimensional remote sensing data with genomic markers [20–25].

Drone (also known as UAS/UAV) imagery provides scalable measurements of phenomic features but comes with logistical and processing challenges. Different flight schedules across environments prevent direct comparisons of date-specific and growth-aligned phenomic measurements. Functional data analysis (FDA) addresses this problem by treating the complete functional trajectory as the unit of analysis rather than an individual time point or scalar summary of a time window [26–28]. Functional principal component analysis (FPCA), a popular FDA tool, has previously led to constructing multi-environment phenomic relatedness matrices using vegetation index (VI) trajectories captured on different flight dates [29]. Functional principal components (functional PCs) provide heritable summaries of temporal traits for biological modeling [30, 31].

Comprising 356 temporal drone flights across 19 year–location combinations, the 2020-2021 Genomes to Fields (G2F) experiments (released publicly via Purdue University’s Data to Science, D2S) represent the largest public field image reference data set to date. This complements preexisting genotype, phenotype, and weather data released every two years through G2F [32, 33]. Differences in environmental conditions and flight timing complicated the integration of drone observations from the same G2F genetic material for modeling and genetic inference. The resource released here enables comparisons of how temporal phenomic data are integrated and interpreted across environments.

As high-dimensional, multimodal data sets become ubiquitous across research programs, analytical approaches for synthesizing input streams without overfitting are increasingly needed. Using this new public data set, we first developed an FPCA-based framework for irregularly sampled phenomic and enviromic trajectories collected across 19 year–locations in the 2020-2021 G2F experiments. Because drone flights cannot be scheduled at synchronous developmental stages across environments, we investigated whether indexing VI and weather trajectories by the physiologically informed time scale of accumulated growing degree days (AGDD) better revealed temporal variation versus the commonly used calendar day scale. We then tested whether functional representations contributed information to grain yield prediction when combined with genomic and enviromic information. Two complementary approaches to temporal tabular data, kernel-based regression and transformer neural processes, were evaluated. Finally, we mapped VI functional PC scores to identify genomic regions associated with temporal spectral variation across environments, linking predictive features to genetic variation. Together, the resource and analyses connect irregular field phenotyping to yield prediction and genetic variation in temporal canopy traits.

## 2 Results

### 2.1 Heritable variation in yield and temporal vegetation indices

Grain yield and vegetation indices showed heritable variation across 19 environments. Heritability varied by environment and drone flight. The normalized green-red difference index (NGRDI), derived from red-green-blue imagery, was used for kernel-based prediction and mapping. The highest-heritability NGRDI values across environments were observed during late vegetative stages, flowering, and senescence (e.g., reaching *H*^2^ = 0.917 for Michigan [MIH1.2020], 57 days after planting [DAP]), with heritability ranging between *H*^2^ = 0.000-0.917 across all environments and flights (Table S6). NGRDI heritability estimates of zero occurred in the earliest or latest flights in environments MNH1.2021, MOH1.2020, TXH1/2/3.2021, WIH1/3.2021. Genotypic variance and heritability of grain yield also varied by environment and within environments that had multiple trials or stress conditions (Figure 1), ranging from *H*^2^ = 0.158 (Texas [TXH3.2020], delay-planted for heat) to 0.788 (Nebraska [NEH1.2021]).

**Fig. 1.**
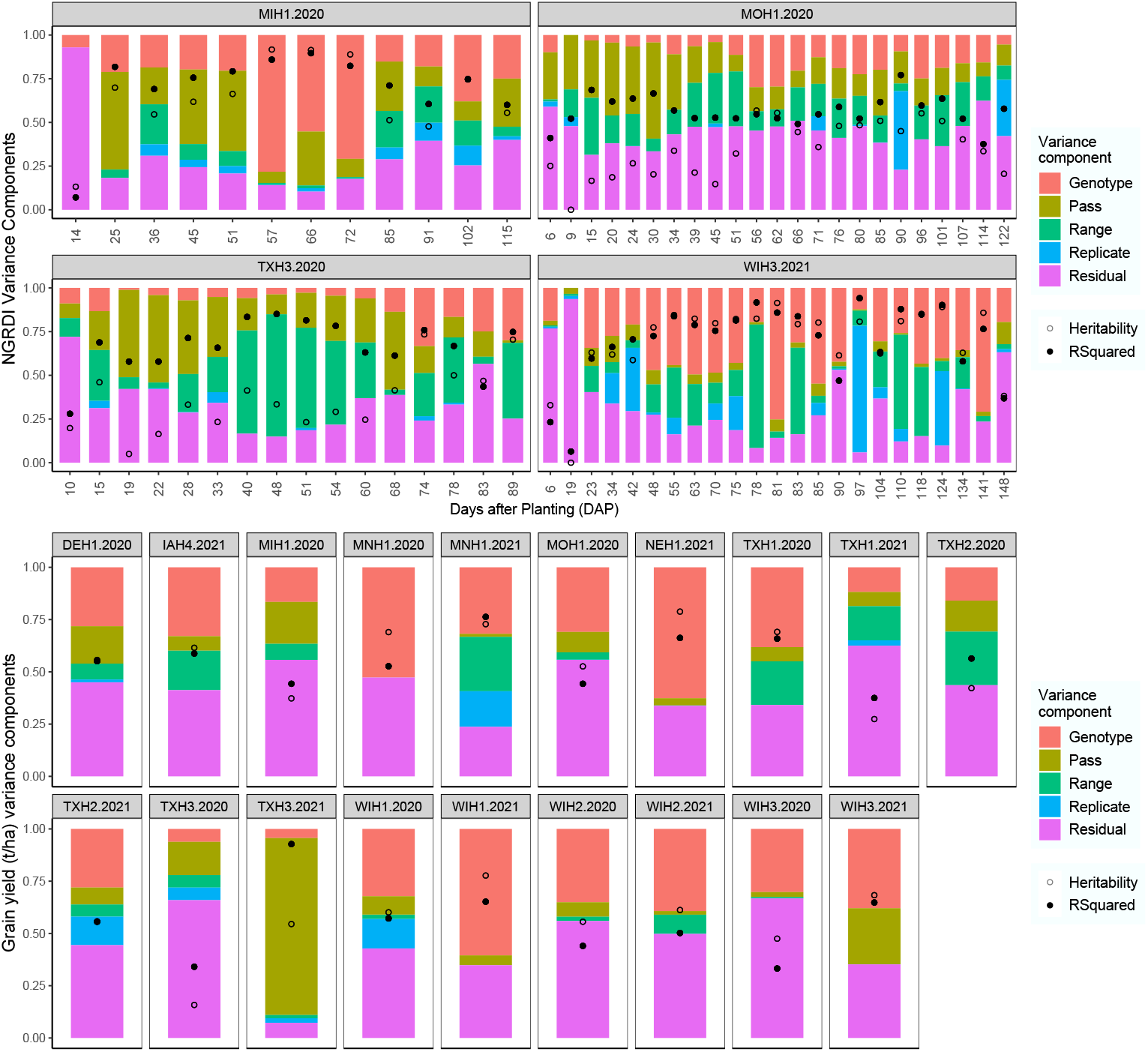
Variance components. (Equation (1)) and heritability values (Equation (2)) for the normalized green-red difference index (NGRDI) in 4 of 19 representative G2F environments (top). Grain yield (t/ha; bottom) variance components and heritability in all 19 environments. For the MOH1.2020 environment, 23 of 45 flights are presented for legibility. Pass and Range are spatial variation model components representing the X and Y axes of the field, respectively.

### 2.2 DAP– and AGDD-indexed FPCA captured complementary temporal variation in VIs

Phenomic approaches have traditionally used DAP, while physiological studies have shown that accumulated growing degree days (AGDD) may better align maize phenology across environmental conditions [34]. FPCA indexed by DAP and AGDD (Figure 2A) characterized variability in VI trajectories throughout the season (Figure 2B). Variation in FPC1 showed contrasting NGRDI values between 37-125 DAP or 175-1450 AGDD (Figure 2). For DAP, FPC1 explained 48.3% of functional variation–the variation in level and shape of a trait’s trajectory over time–in temporal NGRDI values across 19 G2F environments (FPC2 explained 24.7%, Figure 2C). For AGDD (Supplementary Figure S1), FPC1 explained less variation (35.5%) with more picked up by FPC2 (33.4%). Biplot comparisons of functional PCs revealed environmental clustering based on temporal NGRDI. Higher FPC1 scores were generally associated with elevated grain yield values (Figure 2D). Among all VI functional PCs, NGRDI PCs were among the highest-correlated with grain yield best linear unbiased estimates (BLUEs) across all environments (DAP FPC1 *r* = 0.55; AGDD FPC3 *r* = 0.60; Table S6). Within-environment NGRDI correlations between yield and functional PCs ranged dramatically (DAP –0.29-0.72; AGDD –0.50-0.64).

**Fig. 2.**
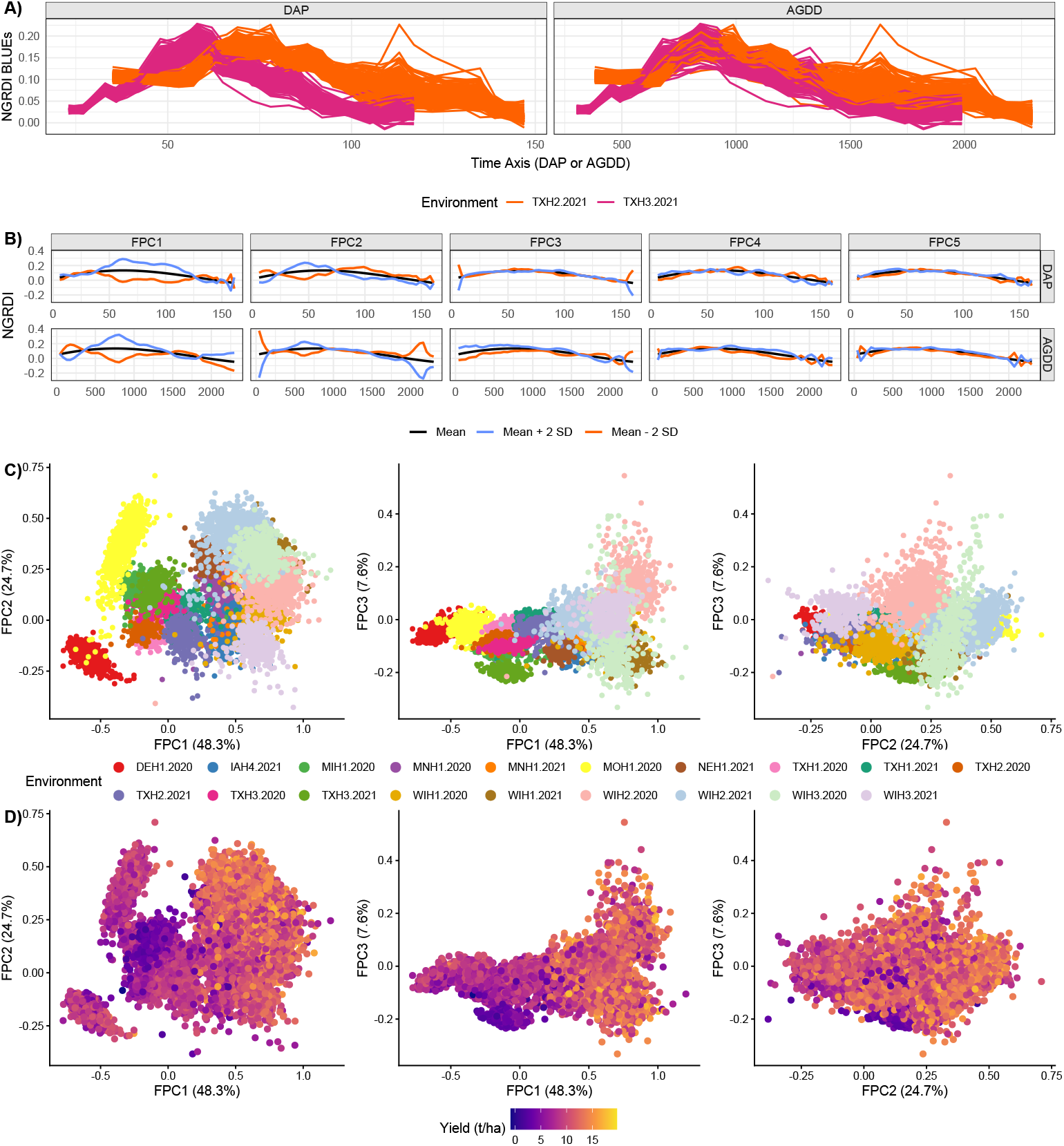
FPCA inputs (A) and results (B-D) from vegetation index data from 19 G2F environments. (A) Temporal registration differences between days after planting (DAP) and accumulated GDD (AGDD) in two Texas environments (TXH2/3.2021; dryland vs. delay-planted heat trials); (B) Perturbations to functional PC scores and their influences on temporal NGRDI values across 19 G2F environments; (C) biplots of FPC1 vs. FPC2/3 and FPC2 vs. FPC3 revealed distinct environmental clusters; (D) biplots overlaid with a grain yield color gradient. Each dot in (C) and (D) relates to one unique combination of a genotype’s values in one of the 19 environments. FPCA results in (C) and (D) are shown for DAP as the time axis (AGDD results shown in Supplementary Figure S1).

Separate FPCA fits to individual weather variables highlighted variability in photothermal ratio (PTR) temporal data and differences in DAP vs. AGDD trajectories (Figure 3A). The first functional PC of the photothermal ratio (PTR; DAP time scale) had both the highest positive correlation and the highest magnitude of correlation with environment-level grain yield (*r* = 0.66), followed by photoperiod FPC1 (N; *r* = 0.62). FPC2 of T2M MIN, the minimum temperature measured at 2 m above the surface, was the most negatively correlated with grain yield (*r* = –0.64), followed by FPC1 of the slope of the saturation vapor pressure curve (*r* = –0.61).

**Fig. 3.**
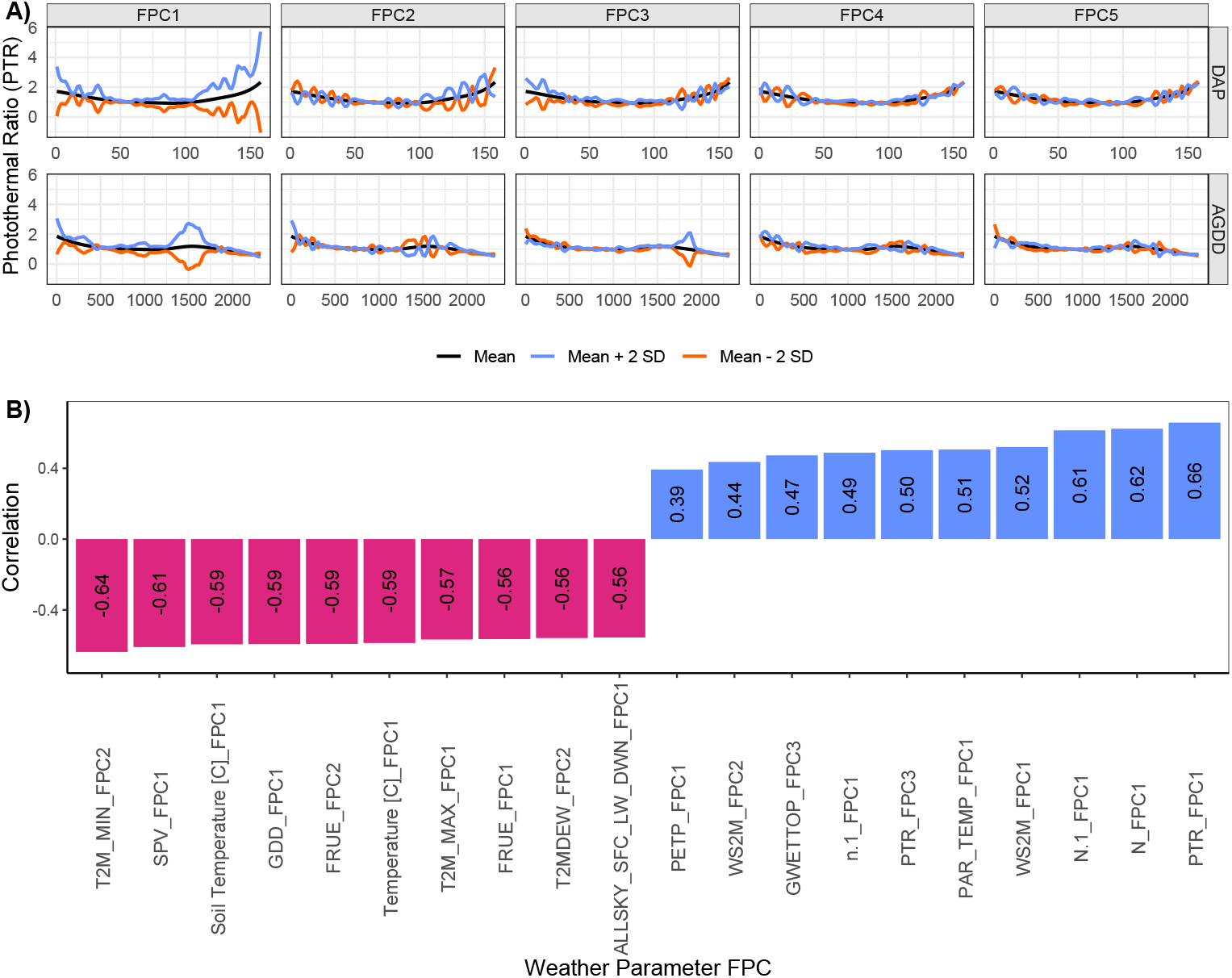
(A) Changes in PTR trajectories when each functional principal component score is shifted by ±2 standard deviations; (B) Correlations of the top 10 most positively– or negatively-correlated environmental parameter FPCs with environment-level yield for 19 G2F environments. Correlations used median grain yield BLUEs from the 10,109 matched records across 19 environments.

### 2.3 Multimodal information improved yield prediction, with performance dependent on the prediction task

The cross-validation (CV) schemes included here—CV2, CV1, CV0, and CV00—refer to prediction scenarios of increasing difficulty. CV2 evaluates models on their ability to reconstruct yield for training hybrids in familiar environments, serving as an in-sample reference. CV1 tests uncharacterized hybrids in familiar environments, CV0 evaluates characterized hybrids in unfamiliar environments, and CV00 evaluates uncharacterized hybrids in unfamiliar environments. A separate analysis, leave-one-environment-out (LOEO) CV omitted all data from one environment at a time from training.

### 2.4 Familiar environments–kernel-based prediction

Prediction performance was strongest when all environments were represented in training (CV2/CV1). Combining genomic and phenomic kernels (M6 and above) improved prediction performance on average by 37.3% (Figure 4A) over each modality alone. Time domain differences (DAP vs. AGDD) were modest. M8.G.P-Int (genomic, phenomic, G×E, P×E) was the top-performing model for CV2 and CV1.

**Fig. 4.**
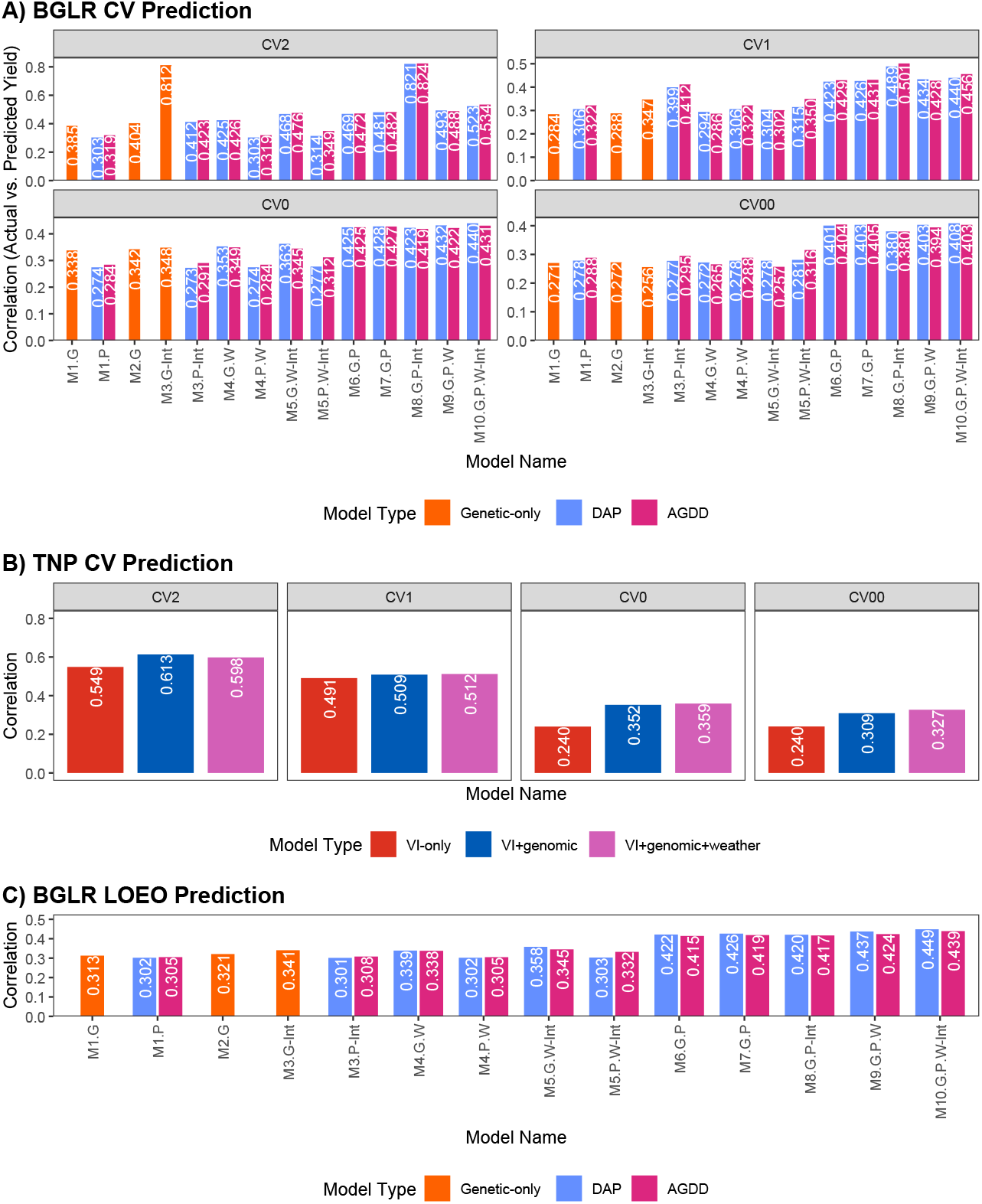
Yield correlations for. (A) 14 kernel models and (B) three transformer neural process (TNP) feature sets across four evaluation schemes, and (C) kernel models under leave-one-environment-out cross-validation. Panels A and B show means over five folds within seed, then over 10 kernel or five TNP seeds. CV2 is an in-sample evaluation; CV1 withholds specific maternal lines, CV0 omits environments, and CV00 omits both lines and environments from training. DAP, days after planting; AGDD, accumulated growing degree days; VI, vegetation indices. Model key: *A*, *D*, *P* and *W* denote additive genomic, dominance genomic, phenomic and weather components; subscripts *E* and *W* denote environment and weather interactions. M1.G: *A*; M1.P: *P*; M2.G: *A* + *D*; M3.G-Int: *A* + *D* + *A_E_* + *D_E_*; M3.P-Int: *P* + *P_E_*; M4.G.W: *A* + *D* + *W*; M4.P.W: *P* + *W*; M5.G.W-Int: *A* + *D* + *W* + *A_W_* + *D_W_*; M5.P.W-Int: *P* + *W* + *P_W_*; M6.G.P: *A* + *P*; M7.G.P: *A* + *D* + *P*; M8.G.P-Int: *A*+*D*+*P* +*A_E_* +*D_E_* +*P_E_*; M9.G.P.W: *A*+*D*+*P* +*W*; M10.G.P.W-Int: *A*+*D*+*P* +*W* +*A_W_* +*D_W_* +*P_W_*. Full model definitions appear in Supplementary Tables S3 and S4.

### 2.5 Unfamiliar environments–kernel-based prediction

Omitting an entire environment from training reduced prediction accuracy for that environment (CV0), especially when the target hybrids were also absent from training (CV00). Combining genomic and phenomic kernels improved prediction performance on average by 39.4% (Figure 4A) over each individually. The PTR weather kernel did not consistently improve prediction performance. DAP and AGDD performed similarly for M10.G.P.W-Int (hereafter M10), which combines genomic, phenomic, and weather effects with genomic *×* weather and phenomic *×* weather interactions (CV0: DAP *r* = 0.440; AGDD *r* = 0.431).

### 2.6 TNP analysis of raw, irregular temporal trajectories

The transformer neural process (TNP) achieved a slightly higher correlation in CV1 and lower correlations than the best kernel-based models in CV2, CV0, and CV00. The top-performing TNP models (defined as the highest-correlation models) for CV2, CV1, CV0, and CV00 performed –25.6%, +2.1%, –18.4%, and –20.0% (positive is better), respectively, relative to their kernel-based counterparts, with corresponding RMSE values +32.4%, –0.4%, +0.2%, and +0.4% (negative is better). In paired comparisons, the full-modality TNP had lower pooled RMSE than M10 in CV1 and CV2. In CV0 and CV00, it had lower correlations but lower RMSE than AGDD M10, while RMSE was slightly higher than DAP M10 (Supplementary Figure S3). Within CV0 and CV00, relative performance also varied among environments (Supplementary Figure S2). RMSE differences depended on whether errors were pooled across all scored predictions or the 19 environment-specific RMSEs were averaged equally (Supplementary Figure S4). Weather observations were limited to 19 distinct whole-environment weather samples. Learning representations that generalize to unseen environments is difficult even for low-capacity models, especially flexible TNP models.

### 2.7 Leave-one-environment-out–kernel-based prediction

LOEO trained models on all available yield records from 18 environments and evaluated all hybrids in the remaining environment. CV0 (Section 2.5) additionally withheld one maternal line fold from training and evaluated only hybrids from the four training folds, differentiating it from the LOEO scheme. M10 achieved the highest LOEO correlation using DAP (*r* = 0.449, RMSE 2.55 t/ha). With AGDD, its correlation was similar (*r* = 0.439, RMSE 2.77 t/ha). Genetic-only models behaved as expected, with marginal improvements seen when modeling dominance and G×E effects in addition to additive effects. M5.G.W-Int had slightly higher correlations than M3.G-Int, with lower RMSE under DAP but higher RMSE under AGDD. Notably, models incorporating both genomic and phenomic effects (M6 and above) outperformed genomic/genomic+weather or phenomic/phenomic+weather models by 33.0% (average *r* = 0.321 vs. 0.427; Figure 4C).

### 2.8 QTL mapping of functional PCs uncovered candidate loci associated with phenomic traits consistently across environments

Identifying a genetic basis for plant temporal behavior using dense multimodal data can improve biological understanding and ultimately crop performance. Quantitative trait locus (QTL) mapping of NGRDI functional PC scores identified 167 significant peaks across analyses (Table S6), with 71 and 96 discovered for DAP and AGDD, respectively (Figure 5A). QTL on chromosomes 3 (113-185 Mb) and 7 (127-138 Mb) had high concordance across environments (Figure 5B, C). Linkage disequilibrium blocks were small and localized in these regions (Figure 5D), nominating targets for finer mapping.

**Fig. 5.**
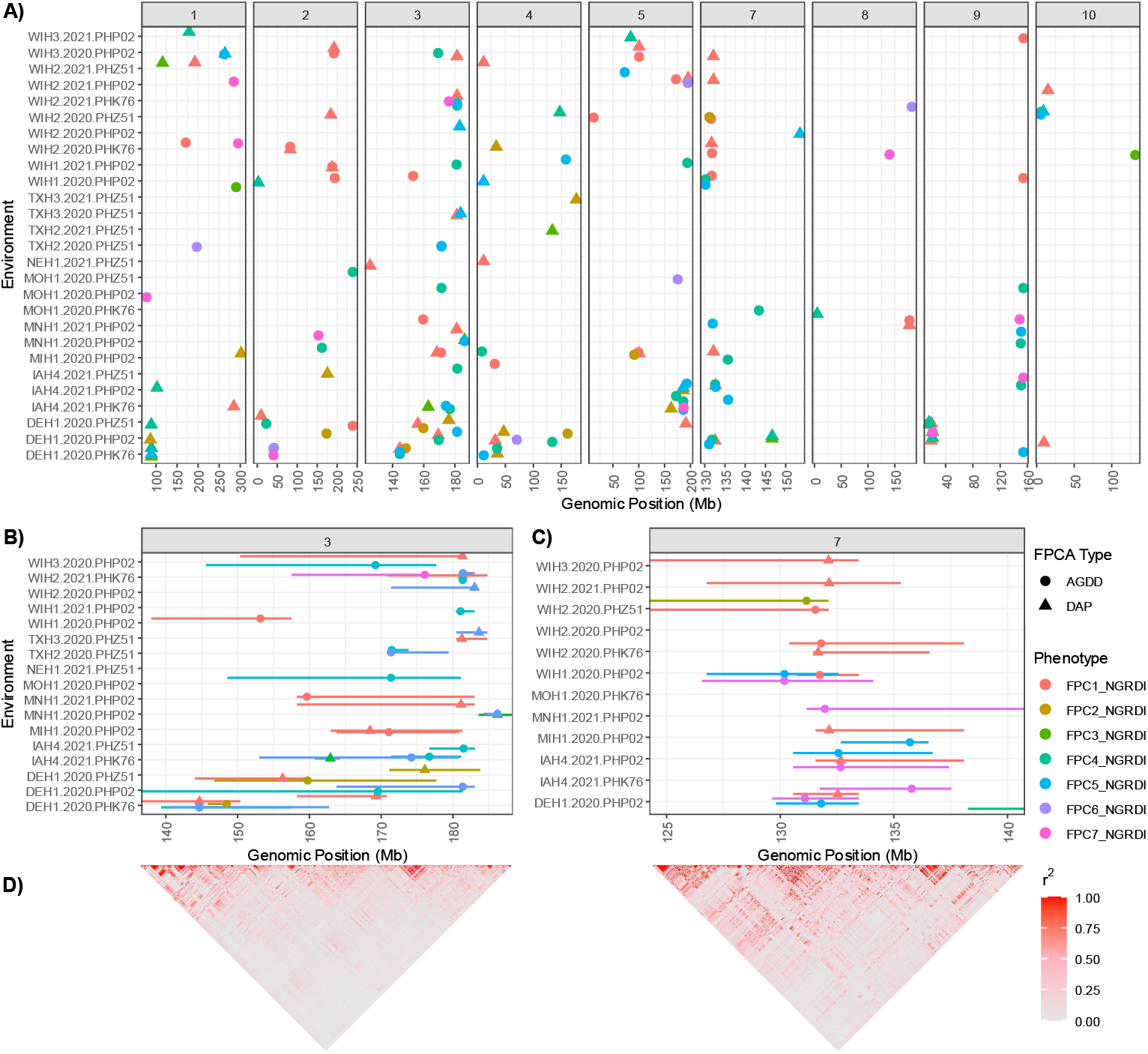
QTL mapping results for each combination of tester and environment of NGRDI FPCs as the traits of interest. (A) 167 candidate loci were discovered across both FPCA methods, with 71 and 96 candidates found for DAP and AGDD FPCA, respectively. Magnification of chromosomes 3 (B) and 7 (C) indicate high concordance of these QTL across environments and testers and low linkage disequilibrium (D) in these regions.

Consistent significant QTL peaks for NGRDI FPC1 on chromosome 7 occurred in nine instances (unique combinations of environment, tester, and FPCA type; Figure 6A top), with multi-parent advanced generation intercross (MAGIC) founders contributing variable BLUP effects (Figure 6A bottom). Gene annotations for the chromosome 7 QTL interval are summarized in Supplementary Table S5. The allele contributed by NKH8431 exerted a strong positive effect on NGRDI FPC1 scores (Figure 6A bottom), associated with high actual NGRDI values immediately before flowering onset and ending at time points that varied by environment (Figure 6B). Alleles contributed by PHJ40 negatively impacted NGRDI FPC1 scores in all environment/tester combinations. MAGIC founder allele effects highlight genetic contributions to variation in NGRDI trajectories summarized by FPC1 (Figure 2B).

**Fig. 6.**
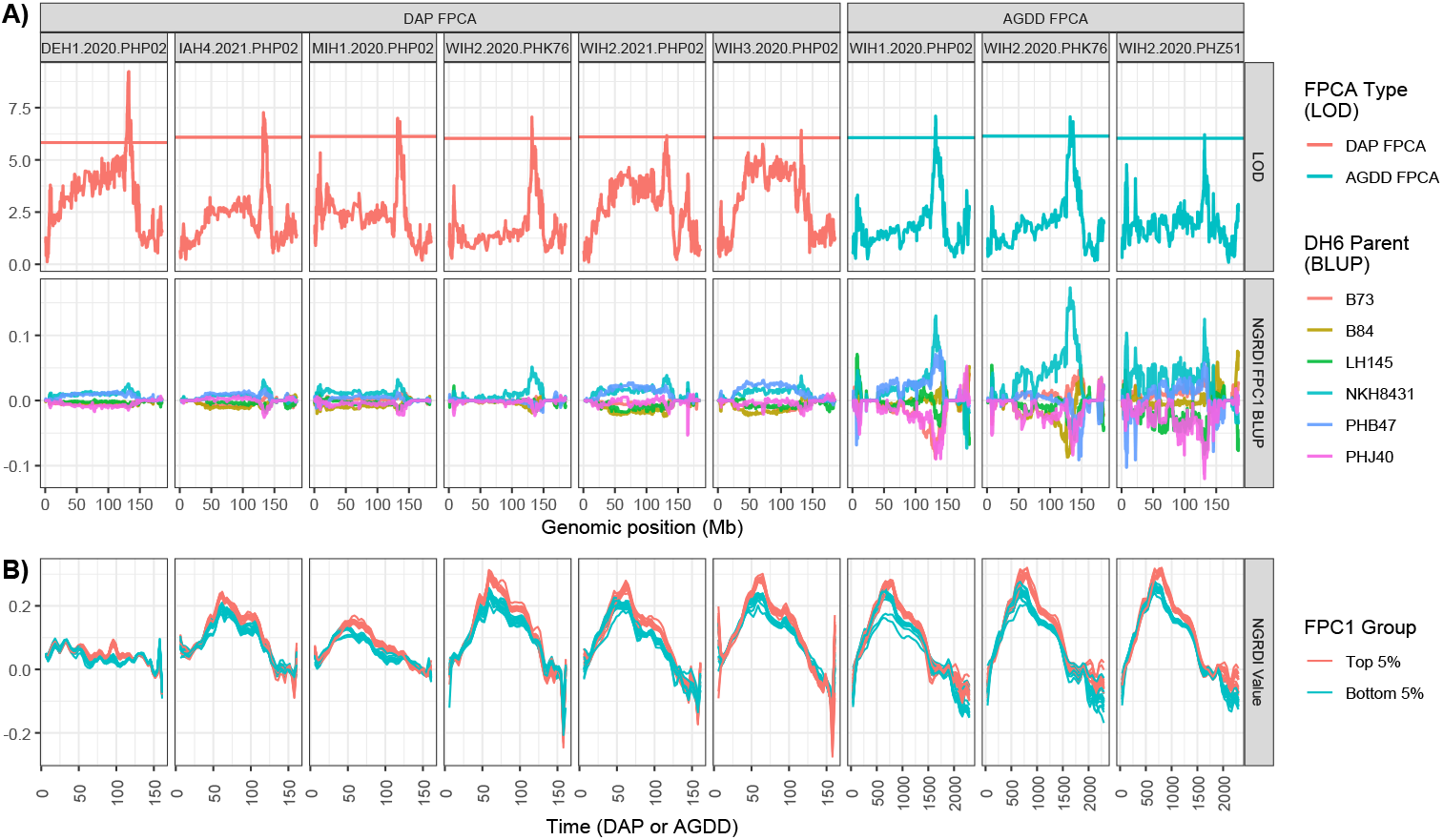
Chromosome 7 QTL results from treating NGRDI FPC1 within environments as a quantitative trait. (A) Top: Loci were deemed significant after surpassing an empirically-defined threshold LOD; Bottom: MAGIC founder lines color coded indicating contributions of their respective alleles. (B) NGRDI values for hybrids with top and bottom 5% of FPC1 scores within each respective tester/environment combination showing temporal divergence of phenotypes.

## 3 Discussion

Extensive temporal drone and weather data are now practical to collect in field experiments across environments to complement conventional genomic data. Combined with conventional genomic prediction methods, multimodal combinations of data types substantially enhanced grain yield prediction across years and locations in this study. Weather data only marginally improved descriptive or predictive power, consistent with previous reports [35–38], though representing weather covariates as functional PCs is novel and open to further study. Methodology has only begun to be developed for integrating these data modalities, especially using phenomic components across environments. FPCA allowed a common representation of irregularly captured vegetation index (VI) trajectories between environments. Phenomic kernels built from functional PCs complemented genomic information in kernel-based prediction. A novel transformer neural process (TNP) model served as a complementary approach that learned directly from raw, irregularly spaced drone observations. Whether larger and more diverse training sets will realize the expected benefits of TNP remains to be tested.

Days after planting (DAP) has been used to index temporal maize phenotypes [39–41], but does not account for differences in heat accumulation across environments. Indexing temporal data by accumulated growing degree days (AGDD), a physiological measure of development, was hypothesized to reduce variation caused by differences in planting dates, weather, and heat accumulation across G2F locations. AGDD resulted in a form of biological curve registration, better aligning observed phenomena to maize developmental time (Figure 2A). Contrary to our hypothesis, AGDD did not consistently improve yield prediction over DAP. This is plausible because thermal time also only approximates maize phenological development, with accuracy depending on the modeling approach. In another study using over 1,000 maize hybrids and more than 50 locations [34], nonlinear and process-based thermal functions predicted time to flowering and the duration from silking to physiological maturity more precisely than simple linear thermal accumulation used here. Similarly, alignment of developmental time to a known phenological landmark, such as panicle emergence in sorghum, better revealed known height loci [30].

Across all modes of data, cross-validation correlations were highest in the in-sample CV2 scheme, followed by CV1, CV0, and CV00 (Figure 4). This CV performance ranking is consistent with the 2014-2015 G2F implementation of these CV schemes [35]. Aligned with a previous analysis of a separate set of 1,918 G2F hybrids across 65 environments, the best yield prediction results were obtained modeling both additive and dominance relationships [42]. While dominance effects did not play a large role here, phenomic trajectories may have served a similar complementary role. Phenomic data likely captured alternative genetic or G×E variation not fully represented by genomic relationship matrices. However, legitimate concerns exist that phenomic prediction can overestimate accuracy because phenomic data are influenced by non-additive and environmental effects [43]. Corrective measures are described for grounded comparisons [43], but approaches to adjust accuracy values from models that simultaneously use genomic and phenomic components still need development.

Surprisingly, incorporating the top FPCs from many other VIs as predictors did not surpass NGRDI alone, nor did a multivariate FPCA approach [44] simultaneously considering all VIs. The higher performance of a single VI is likely attributable here to using a larger number and diversity of environments. Differences in maize phenology, relative to weather conditions across different environments, may render conflicting information from the same VI given both time domains used here. This was supported by within-environment yield vs. NGRDI functional PC correlations ranging between *r* = –0.29-0.72 for DAP and –0.50-0.64 for AGDD. As environments scale, features beyond simply adding more VIs (e.g., textural or morphological) may be required for improved phenomic characterization.

The TNP results reinforced the value of multimodal information streams while highlighting why neural and kernel approaches are complementary rather than direct competitors. TNPs treat prediction with functional predictors as a sequence modeling problem, using attention to learn relationships between observed context points and unobserved targets [45]. Here, adding genomic marker information produced substantial improvement over VI-only TNP. Like the kernel-based models, contributions of weather information were smaller and variable. The full-modality TNP had a slightly higher correlation in CV1 (0.512 vs. 0.501) but did not exceed other models in the remaining CV schemes. Lower correlations did not always coincide with larger prediction errors, highlighting the importance of considering both correlation and RMSE when comparing models. Similar context dependence has been reported in image-based deep learning studies, with a multimodal wheat model demonstrating improved yield prediction relative to genomic-only baselines [24]. Deep networks and BLUP models with interactions had similar prediction error profiles for G2F data, where no singular data source was sufficient in every setting [46]. The TNP learned from raw DAP-indexed VI and weather sets, while kernel-based models used NGRDI functional PCs and a PTR environmental kernel, meaning the two approaches imposed different representations and inductive assumptions.

QTL mapping analysis was used primarily to determine if common heritable genetic loci were observed across different environments. Functional PCs of NGRDI demonstrated common QTL on chromosomes 3 and 7 and connected major temporal modes of canopy variation to segregating genomic regions. Previous longitudinal phenotyping studies have revealed the time-dependent architecture of QTL [47, 48]. FPCA of time-series sorghum height promoted the discovery of known causal loci relative to individual time points or terminal measures [30], temporal maize NDVI revealed genetic effects that varied across development [39], and high-throughput measurements across 16 developmental stages exposed a dynamic architecture of maize growth that contained stage-specific QTL and QTL hotspots [49]. Drone measurements of plant height have similarly facilitated discovery of loci acting across and at specific stages in maize [48, 50], wheat [51, 52], and sorghum [53]. Here, repeated detection of chromosome 3 and 7 regions across tester-environment combinations demonstrated that some environments provided better contrast. These loci will be useful for closer studies of genetic effects on maize temporal architecture through gene expression data or finer mapping.

This resource connects temporal drone phenotyping with multimodal yield prediction and genetic analysis. Expanding beyond the 19 environments sampled over two years, particularly outside Texas and Wisconsin, would broaden environmental coverage and enable tests of predictive ability in future seasons and broader germplasm. The present analyses used entire-season VI and weather trajectories, so their utility for decisions earlier in the season remains to be established. A next step is to compare TNP and kernel models using only observations available at successive prediction dates, testing how predictive performance changes as new measurements become available.

Temporal phenomics provides predictive features and captures genetic variation. Phenomic selection has previously been proposed as a scalable, indirect prediction strategy [17], and temporal field phenotypes have improved prediction in maize, wheat, and many other crops [24, 29, 40]. The present study adds an unprecedented across-environment comparison of functional time domains, structured prediction kernels, and a neural process, while connecting the same temporal summaries to QTL analysis. Future analyses should focus on more diverse environments, prediction from truncated time-series trajectories, and validation of the chromosome 3 and 7 QTL regions, potentially leading to mechanistic insights about maize temporal architecture across genetic backgrounds.

## 4 Methods

### 4.1 Genomes to Fields (G2F) experimental design

The 2020-2021 G2F experiments tested the performance of the Wisconsin Stiff-Stalk Multi-parent Advanced Generation Intercross (WI-SS-MAGIC) population described in Michel et al., 2022 [54] as hybrids crossed with three locally-adapted testers, namely PHP02 (Northern locations), PHK76 (Midwest/intermediate), and PHZ51 (Southern). WI-SS-MAGIC contains subset populations “A” and “B”, where before doubled haploid induction, “A” individuals underwent four instances of meiosis and B individuals underwent six. Local commercial checks were included in each location and experimental checks (“yellow stripes”) were included in all locations. In total, 19 distinct environments were evaluated (location–treatment–year combinations) and are abbreviated by two-letter state codes followed by “H”, the number of the location or treatment, and ending in the year (e.g., TXH1.2020 indicates Texas, treatment 1 [optimal planting date and irrigated], year 2020). Trials were planted as modified randomized complete block designs, with each trial generally containing two replications of each hybrid (500 plots) planted in two-row plots. Each plot was approximately 6 m in length with 0.76 to 1.83 m alleys between plots. Planting dates, latitudes, longitudes, and elevations are indicated in Table S6.

### 4.2 G2F high-throughput phenotyping and drone data processing

RGB drone data were collected from each of the 19 environments at varying temporal resolutions. Camera types, ground sampling distances, and flight dates (with days after planting) are outlined in Table S6. RGB images were stitched into orthomosaics using Agisoft Metashape Version 2.0.2 (Agisoft LLC, St. Petersburg, Russia). Specific Metashape parameters used are provided in the Supplementary Methods. Plot shape-files were created with UAStools [55] and adjusted manually with QGIS [56]. Pixels with HUE *>* 0 were masked, and each vegetation index was averaged over the remaining non-missing pixels within each plot polygon using FIELDimageR [57]. Plot-level raw phenomic data for each time point are available in Table S6. These data were adjusted for field effects within each flight/environment combination using Equation (1), and the genotype means formed the temporal trajectories.

### 4.3 Enviromic data

Enviromic data were obtained from NASA POWER using EnvRtype [4] and G2F infield weather stations. The beginning of the enviromic data collection window was defined as the date of planting and the final date was defined as the last drone flight date in each environment to ensure synchronization of phenomic and enviromic data. EnvRtype weather variables and descriptions are provided in Table S6. In the processWTH() function, temperature limits for GDD were 8 and 45 *^◦^*C, and optimum temperature parameters for radiation use efficiency were 30 and 37 *^◦^*C. In addition to the default EnvRtype variables, photothermal ratio (PTR) and photothermal time were calculated as the photoperiod divided by the growing degree days (GDD) and GDD multiplied by the photoperiod, respectively [58]. Air and soil temperature (*^◦^*C) were sourced from in-field weather stations [33]. Data cleanup and outlier removal procedures are documented in the EnvRtype script [59] and the cleaned weather data are provided in Table S6. As the enviromic data were analyzed with FPCA before being included in prediction models, missing data or data removed during cleaning were handled by the fdapace package. For TNP models (below), the daily weather series were consumed directly; the few missing daily values that remained after cleaning were carried forward from the previous observed day within each environment, as described in the Supplementary Methods.

### 4.4 Genomic data

Genomic data for G2F hybrids were downloaded from the 2024 G2F Genotype by Environment Prediction Competition CyVerse page [60]. For this study, 1,180 hybrids with genomic and phenomic data (Table S6) were used to subset the full VCF file, which was saved as a HapMap file using TASSEL [61]. These hybrids consisted of a mixture of experimental check hybrids and hybrids created by crossing subsets of 383 unique MAGIC inbreds (females) with three regionally specific testers (males). Of the initial set of 437,214 available markers, 238,782 remained after performing the following filtering procedure: i) retain only biallelic markers; ii) remove any marker in which one or more hybrids have a missing value associated with it; iii) remove markers with a minor allele frequency less than 0.05. Filtering steps were performed in base R and rTASSEL [62]. After filtering, the HapMap file was converted to a dosage matrix.

### 4.5 Phenotypic and phenomic data mixed models

Agronomic phenotypic data, including unique plot IDs, field spatial coordinates (pass – X axis of the field, synonymous with “row”; range – Y axis), genotype names, and plot yield values (t/ha) were obtained from CyVerse [63, 64]. Best linear unbiased estimates (BLUEs) were obtained before downstream analysis [65] using the R package lme4 [66]. One model was fit per environment for yield and per VI–environment–flight combination for vegetation indices.

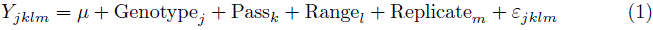

In this equation, *µ* denotes the grand mean, Genotype*_j_* denotes the effect of each genotype *j*, Pass*_k_* denotes the effect of each X coordinate *k* of the field, Range*_l_* denotes the effect of each Y *l* coordinate of the field, Replicate*_m_* denotes the effect of the replicate *m* of each genotype, and *ε_jklm_* denotes the residual error unexplained by the predictors. Pass (*k*) and Range (*l*) numbers vary by environment. The exact numbers of each Genotype, Pass, Range, and Replicate are provided in Table S6.

Genotype, Range, Pass, and Replicate were set as factor variables. To obtain BLUEs, Genotype was specified as a fixed effect while the other effects were specified as random. To compute heritability and variance components, the same models were implemented with all variables (including Genotype) specified as random effects. Entry-mean broad-sense heritability was calculated as:

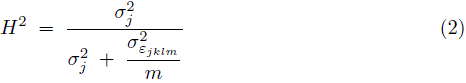

In Equation (2), *m* indicates the number of replicates, 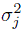 indicates the variance attributed to the Genotype variance component, and 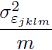 denotes the residual error variance divided by the number of replicates. After the BLUEs analysis, data corresponding to 10,109 genotype/environment combinations were available for downstream analysis (Table S6). This number arose from the overlap between Genotypes with valid values in both phenotypic (e.g., yield) and phenomic (VI) data.

### 4.6 Functional principal component analysis for sparse phenomic and enviromic data

#### 4.6.1 Functional principal component analysis (FPCA)

The FPCA function within fdapace was used as a framework to analyze sparse, nonoverlapping phenomic and enviromic data. The package allowed the use of the native R predict function as a wrapper to project held-out curves onto training basis functions, enabling the implementation of a data leakage prevention strategy. Borrowing notation from Yao et al., 2005 [67], this function implemented the model:

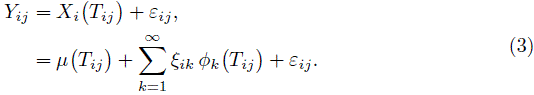

Here *X_i_*(*·*) denotes the latent random function; *T_ij_* specifies the time of observation *j* for trajectory *i*, indexed by DAP or AGDD; *Y_ij_* denotes the *j*th observation of the random function *X_i_*(*·*), where *i* is an index for either phenomic data (*i* = 1, …, 10, 109) or enviromic data (*i* = 1, …, 19); *µ*(*·*) denotes the mean function; *ξ_ik_* symbolizes the *k*th functional principal component (FPC) score associated with *X_i_*(*·*); *ϕ_k_*(*·*) denotes the *k*th eigenfunction; and *ε_ij_* are measurement errors assumed independent of the FPCs. For each VI, trajectory *i* is the series of adjusted values (BLUEs) for one genotype in one environment (10,109 curves total). Each weather variable was the daily series in one environment (19 curves). The two data types were modeled separately. The flexibility of this framework stems from its tolerance for indexing observations both by their individual phenomic or enviromic functions, *i*, and observations, *j*, allowing both sparse and nonoverlapping measurements (as given by *T_ij_*). FPCA for phenomic and enviromic data was performed first by indexing the time domain by days after planting (DAP) and second by accumulated GDD (AGDD). AGDD was calculated as the cumulative sum of daily GDD from EnvRtype within each environment. Expressing time as accumulated heat rather than elapsed days was intended to align phenology across environments. For DAP and AGDD FPCA, the top five and seven FPCs respectively were retained after testing yield prediction performance beginning with one functional PC up to eight. Exploratory yield prediction comparisons identified five DAP and seven AGDD NGRDI components as the smallest sets attaining the best observed performance. These counts were held fixed across validation splits.

#### 4.6.2 Data leakage prevention with fdapace::predict

For each leave-one-environment-out (LOEO) split, FPCA estimated the mean trajectory and eigenfunctions–the main patterns of temporal variation–from the training environments. Functional PC scores for curves from the held-out environment were then estimated using these fixed functions, analogous to applying a new season’s data to a model trained on previous years. This procedure was implemented with VIs and weather data and was repeated for all 19 environments. The models described in Supplementary Table S4 were formed separately for each held-out environment. Prediction results were aggregated across environments as described in Section 4.8.

For CV2, CV1, CV0, and CV00 results, FPCA was performed only with hybrids assigned to the training data set for their respective CV type for VIs. A fold is one of five groups of shared maternal lines. All hybrids with the same maternal parent were assigned to the same fold, regardless of tester or environment. For CV2 and CV1, this meant that hybrids in four out of five folds in all 19 environments were used to generate training basis functions, while all remaining curves were projected onto these bases to obtain predicted functional PC scores. For CV0 and CV00, hybrids in four out of five folds in all except the held-out environment were used to generate the training basis functions, with all remaining curves projected. Full details of these CV schemes are provided in Section 4.8.

### 4.7 Prediction kernels

#### 4.7.1 Genomic relationship matrices

Using the dosage matrix (4.4), additive and dominance genomic relationship matrices (GRMs) were created using the AGHmatrix R package function Gmatrix by specifying method = “VanRaden” and method = “Vitezica”, respectively [68–70]. Each matrix had dimensions 1, 180 *×* 1, 180 (Table S6; 4.4).

The genotype incidence matrix expanded each additive or dominance GRM to the genotype–environment records, yielding 10,109 *×* 10,109 kernels. Genotype-by-environment kernels were obtained by elementwise (Hadamard) multiplication with a matrix indicating whether pairs of records belonged to the same environment (Supplementary Table S3). In Supplementary Table S3, the following notation is adopted: let *Z_g_ ∈* R^1,180^*^×^*^238,782^ denote the SNP marker matrix of all maize hybrids; let *G_a_* and *G_d_* denote the additive and dominance genomic relationship matrices derived from *Z_g_* using the VanRaden [69] and Vitezica [70] methods, respectively; let *Z_P_ ∈* R^10,109^*^×^*^1,180^ denote the genotype incidence matrix; let *Z_E_ ∈* R^10,109^*^×^*^19^ denote the environment incidence matrix; let *X_p_ ∈* R^10,109^*^×b^* denote the phenomic predictor matrix, where *b* is the number of phenomic variables (i.e., the number of retained functional principal components); let *P* = *X_p_X^⊤^/b* denote the corresponding phenomic relationship matrix; and let *w ∈* R^19^*^×^*^1^ denote the vector of PTR FPC1 scores across the 19 G2F environments, with the environment-level weather relationship matrix defined as *W* = *ww^⊤^ ∈* R^19^*^×^*^19^. FPCA bases were estimated from training curves only. NGRDI functional PC scores were then centered and scaled to unit variance within each environment using both training and projected held-out predictor rows before constructing phenomic kernels across the 10,109 genotype–environment records. Genomic kernels received no additional centering or normalization after GRM construction.

#### 4.7.2 Enviromic relationship matrix

The 19 *×* 19 enviromic relationship matrix was formed by multiplication of the 19 *×* 1 matrix of PTR FPC1 with its transpose. To form the 10, 109 *×* 10, 109 matrix used in the prediction models (to replace the block diagonal matrix in Section 4.7.1), matrix multiplication was performed between the 10, 109 *×* 19 environment design matrix (Section 4.7.1), the 19 *×* 19 enviromic relationship matrix, and the transpose of the environment design matrix. Within each LOEO split and each CV0/CV00 held-out environment scenario, weather functional PC scores were standardized using means and standard deviations estimated exclusively from the non-held-out training environments. These training statistics were then applied to all environments, including the held-out environment. Centering and scaling for CV2/1 scenarios was performed with all data since these scenarios did not hold out an environment. PTR FPC1 was selected before cross-validation based on its correlation with yield across all 19 environments, guided by prior evidence of PTR’s predictive utility [58]. Weather kernel comparisons were therefore conditional on this selected descriptor.

### 4.8 Kernel-based yield prediction with genomic, phenomic, and enviromic data

Ten kernels were combined in 14 models. Kernel definitions and distributional assumptions are provided in Supplementary Table S3, and model configurations are outlined in Supplementary Table S4. Models were each specified using the R package BGLR by setting all kernels as RKHS, which specifies a Gaussian process. Sampling settings, prior specifications, and treatment of environment effects are described in Supplementary Methods (Section S1.4). Two types of cross-validation (CV) were implemented: 1) a modified set of the four CV schemes detailed in Jarquín et al. [35] with variations described as follows due to the nature of multiple testers used to create hybrids in the G2F 2020-2021 field trials, and 2) a LOEO CV scheme, where each environment was iteratively removed from training data and all other environments’ data were used to predict grain yield in the held-out environment. The 223 maternal lines (of 383 total; described in Section 4.4) represented in all 19 environments were randomly assigned to five folds for each of 10 randomizations (seeds 1–10). All tester crosses of each maternal line retained their respective folds across environments. CV2/CV1 used four folds for training across all environments, and CV0/CV00 additionally withheld each environment in turn. Hybrids outside this shared maternal set contributed yield observations from training environments but were excluded from calculations of predictive ability in these four CV schemes.

For both Jarquín et al. [35] CV and LOEO CV, prediction ability is reported as the Pearson correlation (*r*) between actual and predicted grain yield values (for the four CVs, details are provided next; in the held-out environment for LOEO). CV2 prediction ability was evaluated as the correlation between actual and predicted yield values of hybrids in the four of five folds included in the training set, simulating the model’s ability to predict performance of previously characterized hybrids in familiar environments. CV2 can be considered an in-sample evaluation and is expected to have the highest model performance metrics. CV1 assessed the correlation between yield values of hybrids in the fold omitted from training in all environments, simulating the ability to predict performance of uncharacterized hybrids in familiar environments. CV0 evaluated the correlation between yield values of hybrids in the four of five folds included in the training set, but only in the held-out environment, simulating the ability to predict performance of characterized hybrids in an unfamiliar environment. The most difficult scheme, CV00, assessed the correlation between yield values of hybrids in the fold omitted from training and in the held-out environment, simulating the model’s ability to predict performance of uncharacterized hybrids in an unfamiliar environment. As outlined in Section 4.6.2, data belonging to the held-out sets (for either the four CVs or LOEO) used projected functional PC scores to prevent data leakage between the training and held-out data sets. A total of 10 randomization seeds (for assigning fold numbers to females) *×* 5 folds *×* 14 models produced 700 results for CV2/1, and 10 seeds *×* 5 folds *×* 14 models *×* 19 held-out environments produced 13,300 results per time domain for CV0/00. To account for unequal numbers of hybrids grown in each G2F environment, a weighted Pearson correlation (*r_w_*; [71]) was calculated using the following equation:

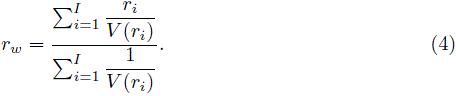

Here, *r_i_* denotes the Pearson correlation between actual and predicted hybrid yield values belonging to environment *i*, and *I* = 19. The sampling variance is

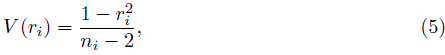

where *n_i_* is the number of paired observed and predicted hybrid yields in each environment/CV scenario. For the four fold-based CV schemes, environment-specific correlations were combined using these weights within each seed–fold combination. Root mean squared error (RMSE) was calculated across all scored records. Both metrics were averaged over folds within each seed and then across seeds. Further aggregation details are provided in Supplementary Methods (Section S1.4). Computations are documented in the accompanying GitHub repository [59].

For models using the phenomic and weather kernels (Supplementary Table S4), prediction results were investigated for kernels created from FPCA using DAP and AGDD as the time domain.

### 4.9 A neural process for multimodal yield prediction

The kernel-based models above rely on FPCA to compress each temporal VI and weather trajectory into a small, fixed-size set of functional PC scores before prediction (Section 4.6). As a complementary, fully data-adaptive alternative, we adopted a deep learning model that operates *directly* on the VI BLUE irregularly spaced time series (at the genotype/entry level), learning its own temporal summaries rather than consuming pre-computed FPCs. The model treats each genotype’s multivariate VI trajectory (the phenomic modality) as an unordered *set* of time-indexed measurements and represents enviromic (weather) data analogously, as a set of daily records; genomic information enters through the genotype’s corresponding rows of the additive and dominance GRMs (Section 4.7.1). Because it consumes sets of (time, value) pairs [72, 73], the model natively accommodates the differing numbers and timings of drone flights and weather records across environments (without interpolation onto a common time grid, binning, or alignment to common time points) and fuses these temporal modalities with the static genomic features in a single end-to-end architecture that produces a probabilistic yield prediction.

The model builds on the neural process family [74], focusing on the subclass of transformer neural processes (TNPs) [45, 75, 76]. Conceptually, the TNP predicts a target genotype’s yield by conditioning on a context set of observed genotypes together with their measured yields, comparing the query genotype against that context to borrow strength from similar observations. This provides an expressive, learned counterpart to the explicit genomic, phenomic, and enviromic relationship matrices used by the kernel-based models (Section 4.7), but one obtained end-to-end and without hand-crafted kernels. The only substantive departure from the standard TNP lies in how each genotype’s heterogeneous VI, weather, and genomic inputs are assembled into the tokens consumed by the transformer backbone [77]; the token construction, network architecture, feature tiers, and training procedure are described in full in the Supplementary Methods, with code in the associated GitHub repository [59].

To isolate the contribution of each modality, the model was evaluated at three nested feature tiers: (i) VI only, (ii) VI + genomic, and (iii) the full VI + genomic + weather configuration. VI and weather values and their time indices were standardized using training set statistics. Models were assessed under the same CV2, CV1, CV0, and CV00 schemes and the same across-environment weighting of Tiezzi et al. [71] used for the kernel-based models (Section 4.8).

### 4.10 Quantitative trait locus (QTL) mapping of phenomic data

QTL mapping was implemented using the qtl2 R package [78] by testing associations between functional PCs of the phenomic data and genomic regions. QTL analyses were performed separately within each of 29 tester–environment combinations. With the same data filtering procedures used to create the genomic prediction kernels (4.4), some hybrids were removed from the analysis based on the thresholds set in Michel et al., 2022 [54], with programmatic implementation details provided in the paper’s GitHub repository [59]. QTL significance thresholds were empirically determined within each tester/environment combination using 1000 permutations of the scan1perm function. QTL effects of loci that exceeded their respective thresholds were estimated with the scan1blup function. The output of scan1blup was passed to find peaks, where using the parameter drop = 1.5, QTL confidence intervals were reported for each putative locus. For regions with high concordance of QTL discovery across environments, the confidence intervals were used as the upper and lower search boundaries when querying MaizeMine v1.5 [79] using its Python application programming interface (API). MaizeMine returned gene descriptions and Gene Ontology (GO) terms.

To calculate linkage disequilibrium within the high-concordance regions, the QTL mapping genotypic data were imported and converted to a genomic data structure (GDS) file compatible with the SNPRelate R package [80]. The snpgdsLDMat function was used to implement the calculations, setting method = “corr” and slide = –1. Full details are provided in the GitHub repository [59].

### 4.11 Data availability

Processed and unprocessed drone imagery, unique to this study, is available from Data 2 Science (D2S). Geospatial data DOIs are listed in this study’s GitHub and in Table S7. Basic information about the G2F field experimental design is available at the G2F website. Genotyping data are available from CyVerse. Raw data files are available either as Supplementary Files (Table S6) or details for how to access them are provided in the repository documentation if other open source data sets were used. All code used to generate results and make figures in this study is available at the associated GitHub repository [59].

## 5 Acknowledgements

The authors are grateful for the efforts of numerous individuals across the G2F consortium: Amanda Gilbert, Dorothy Sweet, Lindsey Newton, Erin Bunting, and Robert Goodwin (Michigan State University); Colby Bass (Texas A&M University); graduate, undergraduate, and high school employees of the Texas A&M Quantitative Genetics and Maize Breeding Program and all G2F institutions. Portions of this research were conducted with the advanced computing resources and consultation provided by Texas A&M High Performance Research Computing. Mention of trade names or commercial products in this publication is solely for the purpose of providing specific information and does not imply recommendation or endorsement by the U.S. Department of Agriculture. USDA is an equal opportunity provider and Employer. Any opinions, findings, and conclusions or recommendations expressed in this material are those of the authors(s) and do not necessarily reflect the views of the National Science Foundation.

### 5.1 Declaration of generative AI and AI-assisted technologies

While preparing this article, author AJD used GPT versions 4.0-6.0 via their web interfaces and the Codex desktop application for production of scripts related to parallelization of prediction and QTL mapping tasks, modifying R plots, organizing the GitHub repository, and rewriting sentences for brevity and clarity. After using these tools, the author reviewed and edited the content to ensure accuracy. The author accepts full responsibility for the content of the article and GitHub repository.

