## Supplementary Materials for "An open field phenomics resource for multimodal maize yield prediction across divergent environments"

### Supplementary Information

#### S1 Supplementary Methods

##### S1.1 Set-Based Deep Learning Model: Implementation Details

This section documents the transformer neural process (TNP) introduced at a high level in the main text (Methods, "A Neural Process Model for Multimodal Yield Prediction"): the data representation it consumes, the three model variants that define the feature tiers, and the training, validation, and testing protocol under which every reported number was produced. The model predicts grain yield directly from the irregularly sampled vegetation index (VI) and weather time series together with static genomic features, and is trained end-to-end by maximizing the Gaussian likelihood of the observed yields. The full source and configuration files are in the associated GitHub repository [1].

###### S1.1.1 Data representation and preprocessing

The modeling unit is a pedigree/environment combination: 10,109 records across the 19 environments (year/location combinations, e.g., TXH1.2020), each comprising a variable-length, irregularly timed VI trajectory paired with a scalar grain-yield BLUE. Yields and VIs are the same BLUEs used by the kernel-based models (main text, Eq. 1). Each record enters the model through up to three modalities, depending on the variant (Section S1.1.5).

**Vegetation indices (phenomic).** A record's VI measurements are treated as an unordered set of time-indexed elements $\{(t_j, \mathbf{v}_j)\}_{j=1}^{T}$, where $t_j$ is the days after planting (DAP) of the $j$th drone flight and $\mathbf{v}_j \in \mathbb{R}^{37}$ collects the 37 VI BLUEs measured at that flight. The number and timing of flights vary freely across environments and no interpolation to a common grid, binning, or temporal alignment across records is performed: the set representation consumes the raw, unaligned trajectory directly [2, 3]. Where a pedigree has isolated missing VI values at an otherwise observed flight, they are filled before modeling by linear interpolation in DAP using only the other flights of that same vegetation index; outside the observed range the value is held constant at the nearest observed endpoint rather than linearly extrapolated, so no fabricated trend is presented to the encoder as if measured.

**Weather (enviromic).** Weather is an environment-level daily series: one record per calendar day of the season for each of the 19 environments, over 42 daily channels, namely every column of the cleaned `EnvRtype` table (main text, "Enviromic Data"; [4]) other than the environment identifier, the DAP index, and the calendar date. Besides the weather variables proper, this set includes four site and calendar descriptors (latitude, longitude, day of year, and a days-from-start counter equal to DAP) and a few duplicated columns; it was used as released without further curation. Like the VI trajectory, each record's weather series is represented as a set $\{(\tau_k, \mathbf{w}_k)\}$ of daily observations indexed by DAP, and all pedigrees within an environment share that environment's series. There are therefore only 19 distinct weather series in the entire data set, which is the motivation for the deliberately small and heavily regularized weather encoder described in Section S1.1.2. Two of the 42 channels, PTR and PAR_TEMP, carry a small number of missing daily values in the cleaned table. Because the weather encoder consumes a complete daily grid, these cells are filled before modeling by carrying the previous observed day's value forward within the same environment; a gap at the start of an environment's series is filled with the first observed value. No other imputation is applied to the weather series.

**Genomics.** Genomic information enters as the pedigree's own rows of the additive and dominance genomic relationship matrices (GRMs), $G_a$ and $G_d$ in the notation of the main text ("Genomic Relationship Matrices"). Both are built from the filtered SNP dosage matrix $M \in \{0,1,2\}^{n\times m}$ (main text, "Genomic Data"; denoted $Z_g$ there, with $n$ = 1,180 genotyped pedigrees and $m$ = 238,782 markers) exactly as for the kernel-based models. Writing $p_j$ for the allele frequency of marker $j$, computed as half the mean dosage of column $j$ over the genotyped pedigrees, the additive GRM is

$$G_a = \frac{ZZ^\top}{2\sum_{j=1}^{m} p_j\,(1-p_j)}, \qquad Z_{ij} = M_{ij} - 2p_j, \tag{S1}$$

and the dominance GRM is

$$G_d = \frac{WW^\top}{4\sum_{j=1}^{m} p_j^2\,(1-p_j)^2}, \qquad W_{ij} = \begin{cases} -2p_j^2 & \text{if } M_{ij}=0, \\ 2p_j\,(1-p_j) & \text{if } M_{ij}=1, \\ -2\,(1-p_j)^2 & \text{if } M_{ij}=2. \end{cases} \tag{S2}$$

Both matrices are used uncentered, matching the `BGLR` parameterization of the kernel models. The additive and dominance rows for a pedigree are concatenated into a single static genomic feature vector whose width is twice the number of genotyped pedigrees in the GRM fit set; the genomic branch's input width is filled in automatically from that at model construction time. Because the GRMs depend only on genotype, which is fixed before planting and independent of the yield being predicted, they are fitted over all pedigrees, which is leakage-free for the same reason given for the kernel-based models.

**Standardization.** All inputs are standardized using training-split statistics only. The VI channels and the DAP time index are $z$-scored with means and standard deviations pooled over training-split records; the weather day index and the weather values are $z$-scored with statistics pooled over training-environment records only and then applied to all environments, including a held-out one. All statistics use the unbiased $(n-1)$ estimator. The grain yield target is deliberately left unnormalized, so that RMSE is reported in the original units and is directly comparable to the kernel-based models.

**Time-point augmentation (training only).** To regularize the temporal encoders and expose the model to varying sampling densities, the time points of every modality are randomly subsampled during training. One keep probability per modality is drawn for each task ($p_{\text{VI}} \sim \mathcal{U}[0.75, 1.0)$ for VI and $p_{\text{wthr}} \sim \mathcal{U}[0.5, 1.0)$ for weather), and every time point of every record in the task is then retained independently with that probability, on both the context and the query side, with at least five time points always kept per curve (individual curves lose different points). Weather augmentation is active only in the variant that has a weather branch. No subsampling is applied at validation or test time.

##### S1.1.2 Per-modality encoders and token construction

Each modality is first summarized into a fixed-length vector, and the available summaries are fused into a single token per pedigree/environment record. Unless stated otherwise, every multilayer perceptron (MLP) in the model has one hidden layer whose width equals its output dimension. The VI set is passed through a permutation-invariant transformer set encoder [3]: each $(t_j, \mathbf{v}_j)$ element is embedded by per-axis MLPs, one over the time index and one over the VI vector, whose outputs are concatenated and mapped by a joint MLP to a single element embedding (all three MLPs shared across the set elements), two layers of pre-norm multi-head self-attention [5] let the within-record measurements exchange information, and the resulting element embeddings are pooled into one 64-dimensional vector by pooling-by-multihead-attention (PMA; a single learnable seed query attends over the set) [3]. The environment-level weather set is encoded by a separate, lighter set encoder of the same family: a single MLP embeds the concatenated (day index $\|$ 42 channels) of each daily record, with no separate per-axis encoders, followed by one self-attention layer and PMA to a 32-dimensional vector, with dropout 0.3 on the embedding MLP and the attention layer. Genomic information is passed through an MLP with two hidden layers of width 64 over the concatenated additive and dominance GRM rows. The available modality summaries are concatenated and passed through a fusion MLP to form the feature embedding $\mathbf{x} \in \mathbb{R}^{64}$; this token-construction step (how the heterogeneous VI, weather, and genomic encodings are combined into the tokens consumed by the neural process) is the only substantive adaptation relative to the standard TNP formulation [6, 7, 8].

The observed yield is encoded jointly with a *density* channel that flags whether the yield is present: for a context record the channel is 0 and the true yield is supplied, whereas for a query record the channel is 1 and the yield is replaced by zero [9]. Context and query records pass through the *same* yield encoder and the *same* token-projection MLP, which maps the concatenation of the feature embedding $\mathbf{x}$ and the encoded $(\text{yield}, \text{density})$ pair to the final $d_{\text{model}} = 64$-dimensional token; context and query tokens therefore differ only in the yield/density they carry, with no separately learned placeholder for missing yields.

##### S1.1.3 Context–query prediction

Following the neural process framework, records are partitioned into a *context set* of observed records and a *query set* of records whose yields are predicted. A cross-sample transformer then refines the query tokens against the context tokens, so that each prediction is formed by comparing the query record against the observed records, a learned counterpart to the fixed genomic, phenomic, and enviromic kernel similarities used by the kernel-based models. To keep this step tractable at the context sizes used here (up to ∼8,000 records), the cross-sample transformer is an induced-set (pseudo-token) attention stack of four pre-norm layers [3, 10, 8, 9]: $K = 256$ learnable inducing points first attend over the context, then the context and query tokens attend over those inducing points, reducing the cost of attending over a context of size $N_c$ from $\mathcal{O}(N_c^2)$ to $\mathcal{O}(K\, N_c)$. The refined query tokens are decoded by an MLP with two hidden layers into the parameters of a Gaussian predictive distribution over yield (Section S1.1.4).

During training the model sees *tasks*, each a context/query partition drawn from the training-visible records. Each optimization step uses a batch of four tasks, and each epoch consists of 64 such batches. One context size $n_c \sim \mathcal{U}\{4096, \ldots, 8192\}$ and one query size $n_q \sim \mathcal{U}\{4, \ldots, 32\}$ (discrete uniform draws with both endpoints inclusive) are drawn per batch and shared by its four tasks (redrawn if $n_c + n_q$ exceeds the pool size); each task then draws $n_c + n_q$ distinct records without replacement and splits them into disjoint context and query sets. When a split's training pool is smaller than these ceilings, each ceiling is lowered to the pool size minus one (each side must leave at least one record for the other), so the same fixed configuration works on every split rather than failing on the smaller ones. These large context sizes train the model in the same regime in which it is deployed. During training, the monitoring context is the training pool with the validation carve removed (Section S1.1.7), and the carved validation records form the query set. At final scoring no carve is applied: the context is the *entire* training-visible pool for the split (every FIT and OBSERVE record, including the maternal lines that were carved out for validation during training), and the scored records form the query set.

##### S1.1.4 Predictive head and training objective

The decoder emits two numbers per query record, mapped by a heteroscedastic Gaussian head to a per-record predictive distribution $N(\mu, \sigma^2)$. The mean is constrained non-negative via a softplus, $\mu = \text{softplus}(\cdot)$, and the standard deviation is $\sigma = 0.99\,\text{softplus}(\cdot) + 0.01$, where the additive floor keeps the predictive variance strictly positive. The model is trained end-to-end to minimize the mean per-record negative Gaussian log-likelihood of the observed query yields.

##### S1.1.5 The three TNP variants (feature tiers)

To isolate the contribution of each modality, three nested variants were trained and evaluated. They are *like-for-like*: the VI encoder, the cross-sample encoder, the decoder, the head, the task sampler, the optimizer, the epoch budget, the validation carve, and the cross-validation splits are byte-identical across them, and they differ only in which modality branches exist and, consequently, in the input width of the fusion MLP. Dropping a modality removes its data processor as well as its sub-encoder, so a dropped modality is never loaded, never standardized, and never augmented.

**VI only.** The set encoder over the raw VI trajectory is the sole branch; the fusion MLP maps its 64-dimensional summary to the token feature space. No weather series and no genomic features are read.

**VI + genomic (weather-free).** Adds the genomic branch: the MLP over the concatenated additive and dominance GRM rows. The fusion MLP takes the concatenated 64+64 dimensional summaries.

**VI + genomic + weather (full).** Adds the weather set encoder, giving a fusion input of $64 + 32 + 64$ dimensions.

Table S1 states exactly what changes between them, and Table S2 gives the shared architecture and optimization settings.

##### S1.1.6 Training and optimization

Each model is optimized with AdamW (learning rate $5 \times 10^{-5}$, weight decay 0.01) at a constant learning rate, with gradient-norm clipping at 0.1, for a budget of at most 250 epochs, with early stopping on a carved validation set (Section S1.1.7). Weights are re-seeded immediately before model instantiation so that the initialization depends only on the run's seed and not on how much randomness the data-processing stage consumed.

Table S1: The three TNP variants. Everything not listed (VI encoder, cross-sample encoder, decoder, head, task sampler, optimizer, epoch budget, validation carve, and cross-validation splits) is identical across the three. “Fusion input” is the width of the concatenated modality summaries entering the fusion MLP, whose output is $d_{\text{model}} = 64$ in every case.

| | VI only | VI + genomic | VI + genomic + weather |
|---|---|---|---|
| VI set encoder | yes ($d$=64) | yes ($d$=64) | yes ($d$=64) |
| Genomic MLP branch | — | yes ($d$=64) | yes ($d$=64) |
| Weather set encoder | — | — | yes ($d$=32) |
| Fusion input width | 64 | 128 | 160 |
| Weather data loaded | no | no | yes |
| Genomic data loaded | no | yes | yes |
| VI time-point augment. | $\mathcal{U}[0.75, 1.0)$ | $\mathcal{U}[0.75, 1.0)$ | $\mathcal{U}[0.75, 1.0)$ |
| Weather time-point augment. | — | — | $\mathcal{U}[0.5, 1.0)$ |

**Numerical-stability guards.** Three guards prevent a single non-finite value from ending a run: a forward pass that produces a non-finite distribution parameter, a finite forward pass that nonetheless yields a non-finite loss, and a backward pass that produces any non-finite gradient each cause that training batch to be skipped rather than propagated. In the gradient case the accumulated gradients are detached so the optimizer leaves the parameters untouched.

###### S1.1.7 Validation carve and early stopping

**Validation split.** A monitoring validation set is carved from the training pool of each split by holding out whole maternal lines: 30 distinct female genotypes are drawn uniformly at random without replacement from those present in that pool (approximately 9% of it), and every row of a chosen line is moved to validation. The validation genotypes are therefore disjoint from those used for fitting, and the monitor rewards generalization to unseen varieties rather than memorization, deliberately mirroring the regime the CV1/CV00 test quadrants measure. The carve is governed by a validation seed fixed at 0 and decoupled from the model-initialization and fold seeds, so the same carve rule applies across all replicates. During training, validation records are excluded from the context and are never scored, and they are never drawn from the held-out test environment. At final scoring the carve is lifted (Section S1.1.3): the scored CV1, CV0, and CV00 sets are PREDICT records and are unaffected, whereas CV2, whose scored records are training records by design, is scored over the full FIT pool including the carved lines.

**Early stopping and checkpoint selection.** The monitored quantity is the Pearson correlation between observed and predicted yield on the carved validation set (maximized), recomputed at the end of each epoch. The retained checkpoint is overwritten whenever that correlation improves on the best value so far, and selection stops once 25 consecutive validation epochs pass without an improvement exceeding 0.001; the retained weights are therefore the best validation epoch prior to the plateau. All reported TNP results use this checkpoint.

###### S1.1.8 Cross-validation design and role assignment

The TNP is assessed under the same CV2, CV1, CV0, and CV00 schemes used for the kernel-based models (main text, “Kernel-Based Yield Prediction with Genomic, Phenomic, and Enviromic Data”) [11], as two split families that each yield two scored quadrants from one fit. Every record is

Table S2: Architecture and optimization hyperparameters shared by all three TNP variants. Attention blocks are pre-norm; "heads × dim" gives the number of attention heads and the per-head dimension. Branch-specific rows apply only to the variants that carry that branch (Table S1).

| Component | Setting | Value |
|---|---|---|
| MLPs (unless noted) | hidden layers / hidden width | 1 / output dim |
| VI set encoder | embedding dim / layers / heads | 64 / 2 / $4 \times 16$ |
| | pooling | PMA, 1 seed |
| Weather set encoder | embedding dim / layers / heads | 32 / 1 / $2 \times 16$ |
| | dropout | 0.3 |
| | pooling | PMA, 1 seed |
| Genomic encoder | MLP hidden layers / width | 2 / 64 |
| Token fusion | fused feature dim $d_{\text{model}}$ | 64 |
| | yield/density encoder dim | 64 |
| Cross-sample encoder | type | induced-set (pseudo-token) |
| | inducing points $K$ | 256 |
| | layers / model dim | 4 / 64 |
| | heads / feedforward dim | $4 \times 16$ / 64 |
| Decoder | MLP hidden layers / width | 2 / 64 |
| Head | predictive distribution | heteroscedastic Gaussian |
| Optimizer | AdamW (lr / weight decay) | $5 \times 10^{-5}$ / 0.01 |
| LR schedule | | constant |
| Gradient clipping | max norm | 0.1 |
| Epochs (max) | | 250 |
| Early stopping | patience / min improvement | 25 epochs / 0.001 |
| Tasks per batch / batches per epoch | | 4 / 64 |
| Context size $n_c$ / query size $n_q$ | (train, sampled per batch) | $\mathcal{U}\{4096, \ldots, 8192\}$ / $\mathcal{U}\{4, \ldots, 32\}$ |
| Evaluation context | final scoring | full training-visible pool |
| Loss | | Gaussian NLL |

assigned one of three roles: **FIT** records are training-visible with their yield, **OBSERVE** records are training-visible but unscored, and **PREDICT** records have their yields masked and never enter any context, at training or evaluation time. Writing $k$ for the held-out fold, "common" for membership in the shared fold table of maternal lines, and $e$ for the held-out environment:

- **CV2/CV1 (no environment removed).** PREDICT = common $\wedge$ Fold = $k$, scored as **CV1**; FIT = common $\wedge$ Fold $\neq k$, scored as **CV2**; non-common records are OBSERVE. CV2 is an *in-sample* reconstruction probe, exactly as in the kernel-based analysis, its scored records deliberately part of the evaluation context.

- **CV0/CV00 (one environment removed).** PREDICT = (common $\wedge$ Fold = $k$) $\vee$ Env = $e$; FIT = common $\wedge$ Fold $\neq k$ $\wedge$ Env $\neq e$. Scoring is restricted to the held-out environment: **CV0** = common $\wedge$ Fold $\neq k$ and **CV00** = common $\wedge$ Fold = $k$, both within Env = $e$. The remaining masked records stay unscored, so a held-out genotype is unseen everywhere and the held-out environment contributes no training rows.

The CV0/CV00 family is run once per (fold, held-out environment) pair ($5 \times 19 = 95$ runs per variant per replicate); the CV2/CV1 family is run once per fold (5 runs per variant per replicate).

##### S1.1.9 Replication and compute

The complete experiment is five independent replicates of the full grid. Each replicate uses a distinct fold-partition seed (Seeds 1 to 5 of the shared fold table), with the model-initialization seed set equal to it, so replicates differ in both the genotype partition and the network initialization. All randomness in a run is governed by three seeds: the fold-partition seed (the genotype folds), the model seed (weight initialization and task sampling, re-applied immediately before model instantiation; Section S1.1.6), and the validation-carve seed, fixed at 0 for every run (Section S1.1.7), so each of the 1,500 runs is reproducible from its released configuration alone. Each model was trained on a single NVIDIA A100 GPU with 40 GB of memory.

#### S1.2 Orthomosaic processing and plot extraction

RGB images were stitched using Agisoft Metashape version 2.0.2. Photogrammetry settings were generally as follows: 1) RGB images were loaded into a new Metashape project; 2) photo alignment was conducted using referenced preselection with a key point limit of 40,000 and a tie point limit of 4,000; 3) initial bundle adjustment was performed by optimizing the camera calibration with the camera distortion parameters set to f, cx/cy, k1, k2, k3, p1, and p2; 4) if applicable, ground control points were imported via a CSV file and manual tagging of their locations was performed (not all locations recorded ground control points); 5) the dense point cloud was built using default parameters; 6) the digital elevation map was built using default parameters; 7) the orthomosaic was built using the digital elevation map as the surface. For locations that possessed real-time kinematics tagged images, the setting "Load camera location accuracy from XMP metadata" was enabled in Metashape via the Tools > Preferences > Advanced option menu.

Plot polygons were created with `UAStools` and adjusted in QGIS. The package `UAStools` may require an older version of R to avoid dependency issues. For users experiencing issues with this package, the authors recommend using a Python-based version of this package with similar functionality. Scripts to create shapefiles in Python, with a similar command structure to `UAStools`, can be downloaded from this GitHub repository [12]. Pixels with HUE > 0 were masked using `FIELDimageR`, and each plot-level VI was recorded as the mean over the remaining non-masked pixels. The file outlining cameras, flight dates, and ground sampling distances is listed in Table S6.

#### S1.3 FPCA for kernel-based prediction models

FPCA was performed using `fdapace` in sparse data mode, with Gaussian smoothing of the mean and covariance functions and bandwidths selected by generalized cross-validation. Scores were estimated by conditional expectation. Initial decompositions used a 99% cumulative variance criterion, which is distinct from the component counts retained downstream. For phenomic prediction, FPCA bases were fitted to training curves, and held-out curves were projected onto those bases. Exploratory data analysis of NGRDI with FPCA used all available 10,462 hybrid–environment curves, while kernel-based prediction used the 10,109 records retained after matching phenomic, genomic, and yield data.

NGRDI was the sole VI used to construct the phenomic kernels (others were evaluated in initial testing), with five DAP or seven AGDD functional PCs retained. These PC counts were chosen from exploratory yield prediction comparisons as the number of functional PCs after which no further gains in predictive ability were obtained. Thus, model performance is conditional on this exploratory choice.

AGDD was calculated as the cumulative sum of daily GDD within each environment. In data preprocessing for fold-based prediction, observations sharing an AGDD coordinate were averaged within each genotype–environment–VI combination. Duplicate AGDD values were possible if going from one day to the next, the threshold for heat accumulation was not reached, thus contributing zero new GDD. Weather observations sharing an AGDD coordinate were averaged within each environment before FPCA.

##### S1.3.1 Kernel construction and predictor standardization

Phenomic data from training were combined with projected curves. Afterward, centering and scaling to unit variance was implemented separately within each environment using all available predictor rows, including held-out rows. Weather scores were scaled using the 18 training environments in LOEO and CV0/CV00 prediction. In CV2/CV1, scaling was performed using all 19 environments given that data from all environments were included in model training. The phenomic kernel used standardized NGRDI scores, and the weather kernel used PTR FPC1. These kernels and their interactions were formed for all 10,109 genotype–environment records. Genomic relationships received no additional centering or normalization. Held-out yields were masked during model fitting.

#### S1.4 Kernel model fitting

Kernel models were fitted using `BGLR` in R 4.4.2 with 10,000 Gibbs-sampling iterations, a burn-in of 1,000 iterations, and the default thinning interval of five. Kernel effects followed Gaussian priors. Kernel and residual variances used the default scaled inverse chi-square priors with five degrees of freedom. Prior scale parameters were calculated automatically by `BGLR`, with the default prior calibration setting `R2 = 0.5`. Predictions were posterior means.

For the reported DAP and AGDD analyses, the separate categorical environment main effect was omitted during fitting in both the fold-based CV and LOEO workflows. An overall intercept was retained, and weather and environment interaction kernels were included where specified by the model.

Analyses used the custom `BGLR_Mod_4.4.2` environment, which included a patch preserving matrix dimensions when only one kernel eigenvector was retained.

In the main text, the "multimodal" group of models was classified as M6.G.P, M7.G.P, M8.G.P-Int, M9.G.P.W, and M10.G.P.W-Int. The comparison group contained the remaining nine kernel configurations. The two time domain runs for each genomic-only configuration were averaged into one reference value, matching Figure 4. Relative gains were calculated as 100 times the difference between group mean correlations divided by the comparison group mean. CV2/CV1 and CV0/CV00 were combined within their respective comparisons, and LOEO results were summarized separately.

LOEO evaluated each of the 19 environments in turn, fitting each model on the remaining 18 environments. Correlations were combined across held-out environments using inverse variance weights. Pooled RMSE was calculated as the square root of the mean squared error across all scored, held-out hybrid–environment records. LOEO did not use repeated maternal line partitions.

The separate paired supplementary comparisons used the CV2, CV1, CV0, and CV00 schemes, with five maternal line folds for each of seeds 1–5. For CV0 and CV00, Figure S4 compares pooled RMSE with the arithmetic mean of the 19 environment-specific RMSEs. Each metric was calculated within each seed and fold, then averaged over the five folds within each seed. Overall means were calculated across the five seed summaries.

Table S3: Kernel names, descriptions, and assumptions

| Kernel Abbreviation | Description | Normality and distributional assumptions |
|---|---|---|
| $K_{G\_A}$ | Additive genomic kernel (no assumption of $G \times E$) | $K_{G\_A} = Z_P G_a Z_P^\top$; $u_{G\_A} \sim N(\mathbf{0}, K_{G\_A}\sigma_a^2)$ |
| $K_{G\_D}$ | Dominance genomic kernel (no assumption of $G \times E$) | $K_{G\_D} = Z_P G_d Z_P^\top$; $u_{G\_D} \sim N(\mathbf{0}, K_{G\_D}\sigma_d^2)$ |
| $K_{GE\_A}$ | Additive genomic kernel (assuming $G \times E$) | $K_{GE\_A} = (Z_E Z_E^\top) \odot (Z_P G_a Z_P^\top)$; $u_{GE\_A} \sim N(\mathbf{0}, K_{GE\_A}\sigma_{GE\_A}^2)$ |
| $K_{GE\_D}$ | Dominance genomic kernel (assuming $G \times E$) | $K_{GE\_D} = (Z_E Z_E^\top) \odot (Z_P G_d Z_P^\top)$; $u_{GE\_D} \sim N(\mathbf{0}, K_{GE\_D}\sigma_{GE\_D}^2)$ |
| $K_P$ | Phenomic kernel (no assumption of $P \times E$) | $K_P = P = X_p X_p^\top / b$; $u_P \sim N(\mathbf{0}, K_P\sigma_P^2)$ |
| $K_{PE}$ | Phenomic kernel (assuming $P \times E$) | $K_{PE} = (Z_E Z_E^\top) \odot P$; $u_{PE} \sim N(\mathbf{0}, K_{PE}\sigma_{PE}^2)$ |
| $K_W$ | Enviromic (weather) kernel (used for treating W as a main effect) | $W = ww^\top$; $K_W = Z_E W Z_E^\top$; $u_W \sim N(\mathbf{0}, K_W\sigma_W^2)$ |
| $K_{GW\_A}$ | Additive genomic kernel (assuming $G \times W$) | $K_{GW\_A} = (Z_E W Z_E^\top) \odot (Z_P G_a Z_P^\top)$; $u_{GW\_A} \sim N(\mathbf{0}, K_{GW\_A}\sigma_{GW\_A}^2)$ |
| $K_{GW\_D}$ | Dominance genomic kernel (assuming $G \times W$) | $K_{GW\_D} = (Z_E W Z_E^\top) \odot (Z_P G_d Z_P^\top)$; $u_{GW\_D} \sim N(\mathbf{0}, K_{GW\_D}\sigma_{GW\_D}^2)$ |
| $K_{PW}$ | Phenomic kernel (assuming $P \times W$) | $K_{PW} = (Z_E W Z_E^\top) \odot P$; $u_{PW} \sim N(\mathbf{0}, K_{PW}\sigma_{PW}^2)$ |

Table S4: Model configurations. See Table S3 for descriptions

| Model Abbreviation | Prediction Kernels |
|---|---|
| M1.G | $K_{G\_A}$ |
| M1.P | $K_P$ |
| M2.G | $K_{G\_A}$, $K_{G\_D}$ |
| M3.G-Int | $K_{G\_A}$, $K_{G\_D}$, $K_{GE\_A}$, $K_{GE\_D}$ |
| M3.P-Int | $K_P$, $K_{PE}$ |
| M4.G.W | $K_{G\_A}$, $K_{G\_D}$, $K_W$ |
| M4.P.W | $K_P$, $K_W$ |
| M5.G.W-Int | $K_{G\_A}$, $K_{G\_D}$, $K_W$, $K_{GW\_A}$, $K_{GW\_D}$ |
| M5.P.W-Int | $K_P$, $K_W$, $K_{PW}$ |
| M6.G.P | $K_{G\_A}$, $K_P$ |
| M7.G.P | $K_{G\_A}$, $K_{G\_D}$, $K_P$ |
| M8.G.P-Int | $K_{G\_A}$, $K_{G\_D}$, $K_{GE\_A}$, $K_{GE\_D}$, $K_P$, $K_{PE}$ |
| M9.G.P.W | $K_{G\_A}$, $K_{G\_D}$, $K_P$, $K_W$ |
| M10.G.P.W-Int | $K_{G\_A}$, $K_{G\_D}$, $K_P$, $K_W$, $K_{GW\_A}$, $K_{GW\_D}$, $K_{PW}$ |

### S2 Supplementary Results

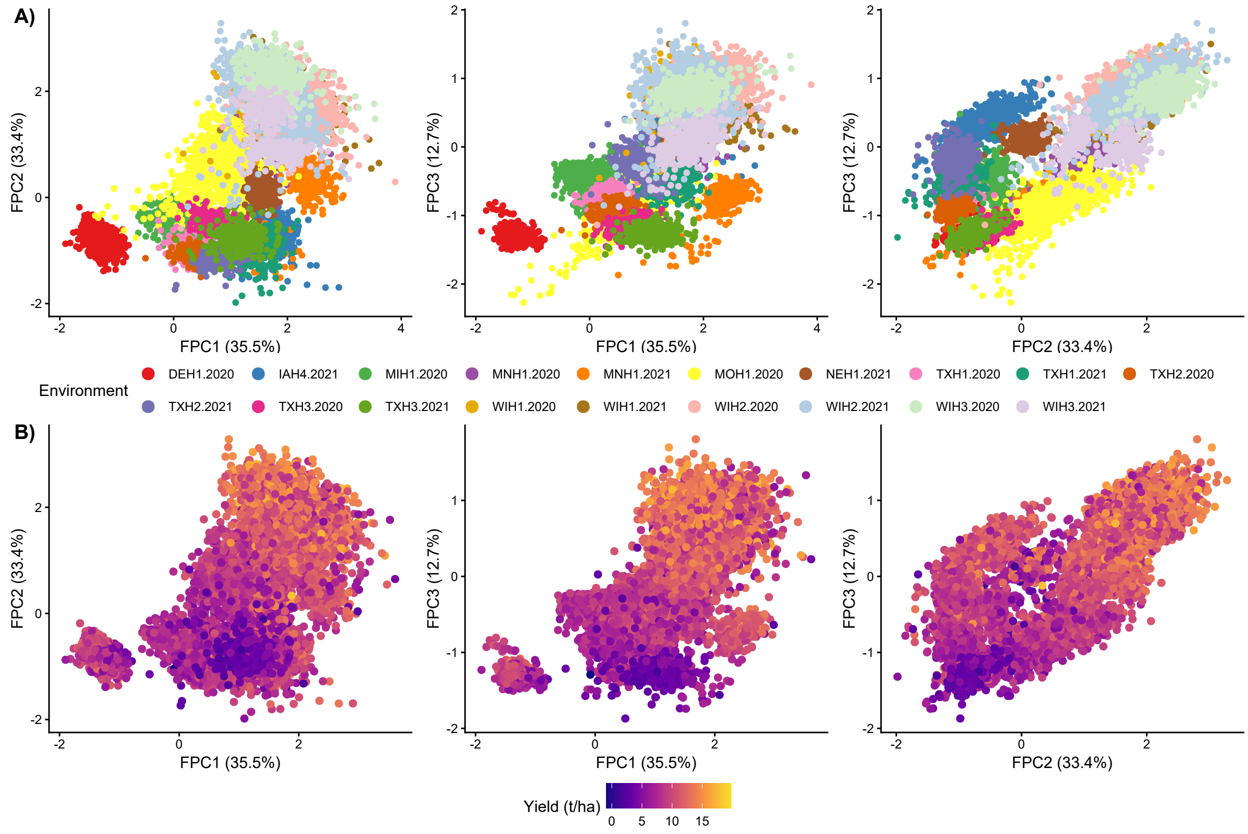


Figure S1: Biplots of the first three functional principal component scores for normalized red green difference index trajectories indexed by accumulated growing degree days. Points represent genotype–environment records and are colored by environment (A) or grain yield (B). Axis percentages indicate the proportion of functional variation explained.

The prediction comparisons in Figures S2 to S4 use M10.G.P.W-Int as a fixed comparator and shared seeds 1-5.

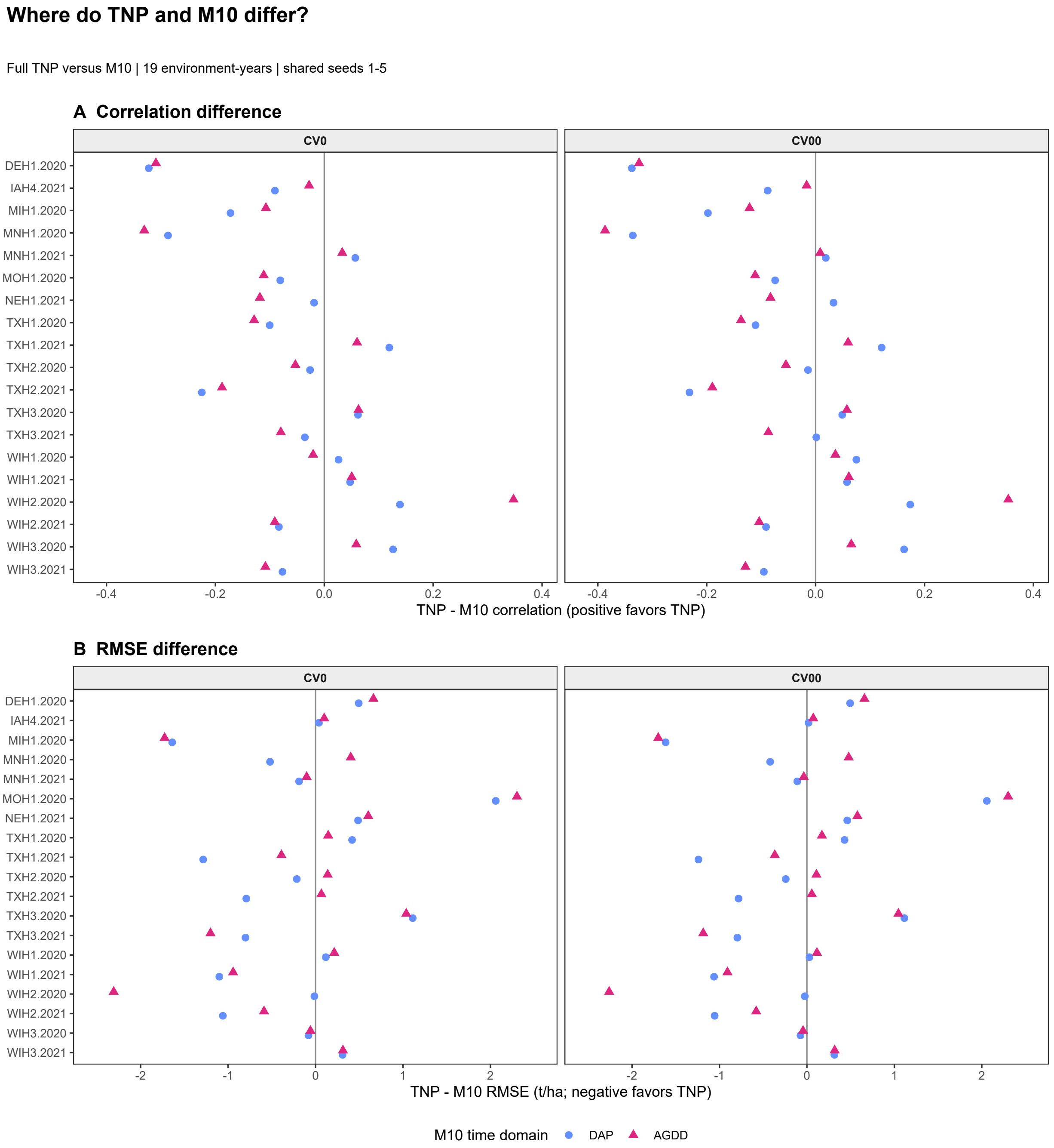


Figure S2: Environment-specific differences between the full transformer neural process (VI+genomic+weather) and M10.G.P.W-Int in Pearson correlation (A) and RMSE (B) under CV0 and CV00. Points show TNP minus M10 for each of the 19 environments after averaging over five folds within each seed and then five shared seeds (1-5). Circles and triangles denote DAP and AGDD M10 results, respectively. The same DAP TNP predictions enter both comparisons. Positive correlation differences and negative RMSE differences favor TNP.

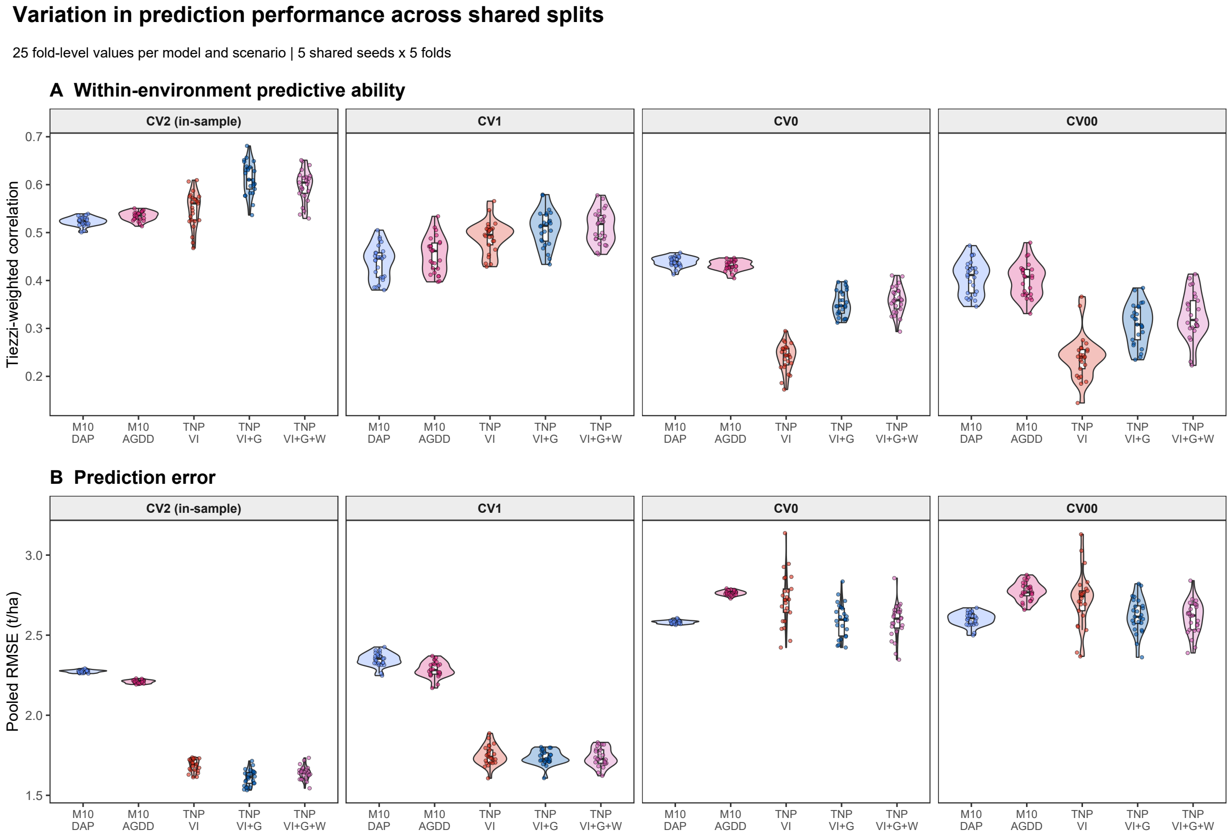


Figure S3: Prediction performance across shared cross-validation splits for M10.G.P.W-Int and three TNP feature sets. Panels show environment-weighted correlations (A) and pooled RMSE (B). Each distribution contains 25 seed-fold values (five shared seeds, five folds). Points show individual values, boxes show medians and interquartile ranges, whiskers extend to observations within 1.5 times that range, and violin widths show smoothed density. These overlapping evaluations describe split variation. VI, G, and W denote vegetation indices, genomic data, and weather, respectively.

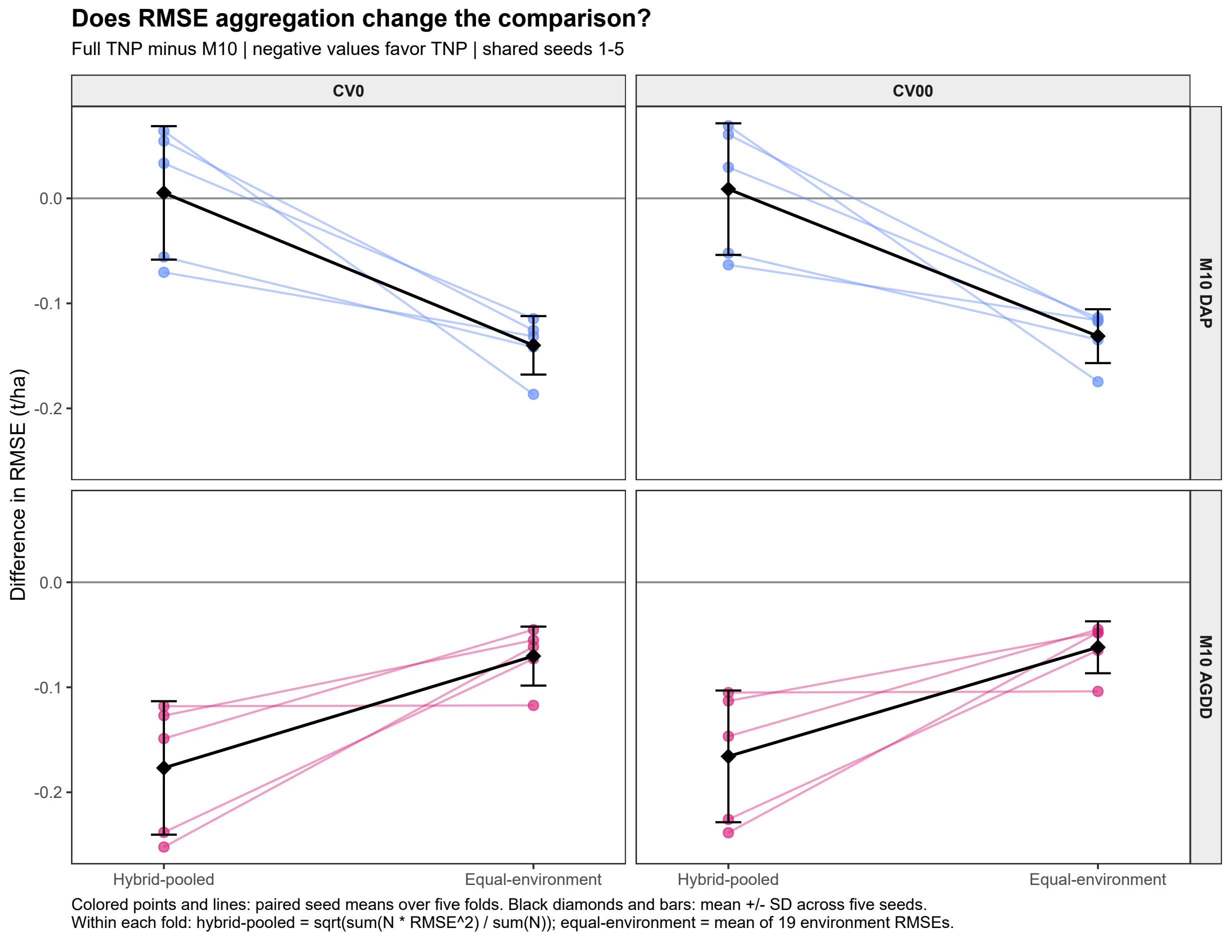


Figure S4: Sensitivity of full TNP minus M10.G.P.W-Int RMSE differences to aggregation under CV0 and CV00. For each combination of seed and fold, pooled RMSE was calculated across all scored predictions, whereas mean environment RMSE was the arithmetic mean of the RMSEs calculated separately for the 19 environments. Colored points and lines connect paired seed summaries averaged over five folds. Black diamonds and bars show the mean and standard deviation across five shared seeds. Negative differences favor TNP. The same DAP TNP predictions are compared with DAP and AGDD M10 versions.

Table S5: Exploratory gene annotations returned by MaizeMine for the chromosome 7 NGRDI FPC1 region. The search window (126,763,800-138,089,853 bp) spans the outer bounds of the clustered QTL intervals in Fig. 5 (main manuscript), excluding the broad WIH2.2020.PHZ51 and WIH3.2020.PHP02 intervals. Records lacking gene symbols or with symbols containing LOC were excluded, and remaining identifiers were grouped by symbol. Listed identifiers include gene models from multiple reference assemblies. Functional descriptions and references provide biological context–causal roles were not explored in this study.

| Symbol | Gene ID(s) | Description | Reference(s) & Context |
|---|---|---|---|
| AT1G16800 | Zm00021ab322740 | p-loop containing nucleoside triphosphate hydrolase superfamily protein | Schenk et al., 2005 [13] (Homolog of *Arabidopsis* Sen1) |
| BADH | Zm00030ab316390, Zm00022ab322390 | Betaine aldehyde dehydrogenase (Fragment) | Fitzgerald et al. [14] (Plant betaine aldehyde dehydrogenase review) |
| BKI1 | Zm00038ab322700 | BRI1 kinase inhibitor 1 | Wang et al., 2017 [15] (Plant architecture regulation in *Arabidopsis*) |
| CSLE6 | Zm00019ab304320 | Cellulose synthase-like protein E6 | Jing et al., 2024 [16] (Copine proteins and brassinosteroid signaling in maize and *Arabidopsis*) |
| CXE2 | Zm00020ab319840, Zm00022ab321080, Zm00023ab324900, Zm00029ab330440, Zm00024ab321920, Zm00032ab329770, Zm00030ab317160, Zm00001eb315150 | Tuliposide A-converting enzyme 2, Tuliposide A-converting enzyme b1 | Chen et al., 2025 [17] (Carboxylesterases in horticultural crops) |
| DMC1 | Zm00026ab320240 | Disrupted meiotic cDNA 1 protein (Fragment) | Gorbunova and Levy, 1999 [18] (Plant DSB repair) |
| MED20 | Zm00042ab329410, Zm00030ab316230, Zm00032ab328970, Zm00001eb314270 | Mediator of RNA polymerase II transcription subunit 20 | Yang et al., 2016 [19] (Plant mediator complexes) |
| OS09G0475500 | Zm00041ab329050 | O-fucosyltransferase family protein | Annotation only–no functional reference supplied |
| RPL2 | Zm00027ab323100 | Ribosomal protein L2 | Adams et al., 2001 [20] (Describes transfer of *rpl2*, a ribonucleoprotein complex responsible for chloroplast protein synthesis, from mitochondria to nucleus in maize) |

Continued on next page

Table S5 continued

| Symbol | Gene ID | Description | Reference(s) & Context |
|---|---|---|---|
| SPL16 | Zm00020ab322440 | SBP-box family member protein (Fragment) | Wei et al., 2024 [21]; Liu et al., 2021 [22]; Wang et al., 2017 [15] (Coordinates short day-induced growth cessation and bud set in *Poplar*) |

#### S3 Supplementary data file inventory

Table S6: Supplementary data and prediction analysis resources. Files accompany the Supplementary Information. Prediction summary code and outputs are available in the repository cited in Code availability.

| Resource | Filename and contents |
|---|---|
| Environment metadata | `Envs_Coordinates.csv`<br>Environment identifiers, coordinates, elevation, state and city. |
| Drone flight metadata | `Drone_Data_Description_V6.xlsx`<br>Flight dates, days after planting, cameras, and ground sampling distances. |
| Cleaned plot data | `G2F_2020_2021_Cleaned_Data.csv`<br>Plot observations, vegetation indices, phenotypes, planting and flight dates, and experimental design identifiers. |
| Phenomic–phenotypic overlap | `Pedigree_Overlap_Phenomic_Phenotypic_BLUEs.csv`<br>Identifiers for 10,109 genotype–environment records shared by the phenomic and phenotypic datasets. |
| Vegetation index and yield variance components | `VarComp_G2F_2020_2021.csv` and `VarComp_Yield_G2F_2020_2021.csv`<br>Variance components for vegetation indices and grain yield. |
| Weather variable definitions | `EnvRType_Weather_Variable_Descriptions.xlsx`<br>Descriptions of EnvRtype weather variables, including units. |
| Air temperature | `Weather_AirTemp_Clean_Tall.csv`<br>Cleaned air temperatures (°C), indexed by environment and days after planting. |
| Soil temperature | `Weather_SoilTemp_Clean_Tall.csv`<br>Cleaned soil temperatures (°C), indexed by environment and days after planting. |
| EnvRtype weather data | `EnvRtype_Weather_Data_Cleaned_V2.csv`<br>Cleaned weather observations and derived variables, indexed by environment and days after planting. |
| Yield BLUEs vs. functional PC correlations | `VI_FPC_Yield_Correlations_Pooled.csv` and `VI_FPC_Yield_Correlations_Within_Environment.csv`<br>Correlation results between yield BLUEs and vegetation index functional PCs. |
| Genomic–phenomic overlap | `Pedigree_Overlap_Genomic_Phenomic.csv`<br>Identifiers for 1,180 hybrids shared by the genomic and phenomic datasets. |
| QTL results | `QTL_Results_Combined_Revised_Intervals.csv`<br>NGRDI FPC results only: 167 significant peaks across analyses, with LOD scores and support intervals. |

Continued on next page

Table S6 (continued)

| Resource | Filename and contents |
| --- | --- |
| Prediction summaries (GitHub) | `Summarize_All_Prediction_Results_V6.R`<br>Code to summarize correlations and RMSE for kernel and TNP models. Primary outputs: `G2F_Final_Prediction_Summary.csv` and `G2F_Final_Prediction_Seed_Summary.csv`. |

le S7: Geospatial data collections hosted on Data 2 Science. The 16 collections cover 19 study environ-
nts. TXH123 combines Texas trials 1–3 within each year, while MOH1.2020 is divided into fields C5a and
[illegible].

| Collection | Data 2 Science | DOI |
| --- | --- | --- |
| DEH1.2020 | Open collection | 10.6084/m9.figshare.33300057 |
| IAH4.2021 | Open collection | 10.6084/m9.figshare.33301599 |
| MIH1.2020 | Open collection | 10.6084/m9.figshare.33301605 |
| MNH1.2020 | Open collection | 10.6084/m9.figshare.33301611 |
| MNH1.2021 | Open collection | 10.6084/m9.figshare.33301701 |
| MOH1.2020.C5a | Open collection | 10.6084/m9.figshare.33301704 |
| MOH1.2020.C5b | Open collection | 10.6084/m9.figshare.33301707 |
| NEH1.2021 | Open collection | 10.6084/m9.figshare.33301713 |
| TXH123.2020 | Open collection | 10.6084/m9.figshare.33301716 |
| TXH123.2021 | Open collection | 10.6084/m9.figshare.33301719 |
| WIH1.2020 | Open collection | 10.6084/m9.figshare.33301749 |
| WIH1.2021 | Open collection | 10.6084/m9.figshare.33301755 |
| WIH2.2020 | Open collection | 10.6084/m9.figshare.33438316 |
| WIH2.2021 | Open collection | 10.6084/m9.figshare.33301758 |
| WIH3.2020 | Open collection | 10.6084/m9.figshare.33301764 |
| WIH3.2021 | Open collection | 10.6084/m9.figshare.33301776 |

[2] Manzil Zaheer, Satwik Kottur, Siamak Ravanbakhsh, Barnabas Poczos, Russ R Salakhutdinov, and Alexander J Smola. Deep sets. *Advances in neural information processing systems*, 30, 2017.

[3] Juho Lee, Yoonho Lee, Jungtaek Kim, Adam Kosiorek, Seungjin Choi, and Yee Whye Teh. Set transformer: A framework for attention-based permutation-invariant neural networks. In *International conference on machine learning*, pages 3744–3753. PMLR, 2019.

[4] Germano Costa-Neto, Giovanni Galli, Humberto Fanelli Carvalho, José Crossa, and Roberto Fritsche-Neto. Envrtype: a software to interplay enviromics and quantitative genomics in agriculture. *G3*, 11(4):jkab040, 2021.

[5] Ashish Vaswani, Noam Shazeer, Niki Parmar, Jakob Uszkoreit, Llion Jones, Aidan N Gomez, Łukasz Kaiser, and Illia Polosukhin. Attention is all you need. *Advances in neural information processing systems*, 30, 2017.

[6] Marta Garnelo, Dan Rosenbaum, Christopher Maddison, Tiago Ramalho, David Saxton, Murray Shanahan, Yee Whye Teh, Danilo Rezende, and SM Ali Eslami. Conditional neural processes. In *International conference on machine learning*, pages 1704–1713. PMLR, 2018.

[7] Tung Nguyen and Aditya Grover. Transformer neural processes: Uncertainty-aware meta learning via sequence modeling. *arXiv preprint arXiv:2207.04179*, 2022.

[8] Leo Feng, Hossein Hajimirsadeghi, Yoshua Bengio, and Mohamed Osama Ahmed. Latent bottlenecked attentive neural processes. *arXiv preprint arXiv:2211.08458*, 2022.

[9] Matthew Ashman, Cristiana Diaconu, Junhyuck Kim, Lakee Sivaraya, Stratis Markou, James Requeima, Wessel P Bruinsma, and Richard E Turner. Translation equivariant transformer neural processes. *arXiv preprint arXiv:2406.12409*, 2024.
